# Differential remodelling in small and large murine airways revealed by novel whole lung airway analysis

**DOI:** 10.1101/2022.01.15.476324

**Authors:** Amanda L Tatler, Christopher J Philp, Michael R Hill, Sam Cox, Andrew Bullock, Anthony Habgood, Alison John, Robert Middlewick, Katherine E Stephenson, Amanda T Goodwin, Charlotte K Billington, Reuben D O’Dea, Bindi S Brook, Simon R Johnson

## Abstract

Airway remodelling occurs in chronic asthma leading to increased airway smooth muscle (ASM) mass and extra-cellular matrix (ECM) deposition. Whilst extensively studied in murine airways; studies report only selected larger airways at one time point meaning the spatial distribution and resolution of remodelling are poorly understood. Here we use a new method allowing comprehensive assessment of the spatial and temporal changes in ASM, ECM and epithelium in large numbers of murine airways after allergen challenge. Using image processing to analyse 20-50 airways from a whole lung section revealed increases in ASM and ECM after allergen challenge were greater in small and large rather than intermediate airways. ASM predominantly accumulated adjacent to the basement membrane whereas ECM was distributed across the airway wall. Epithelial hyperplasia was most marked in small and intermediate airways. Post challenge, ASM changes resolved over seven days whereas ECM and epithelial changes persisted. The new method suggests large and small airways remodel differently and the long-term consequences of airway inflammation may depend more on ECM and epithelial changes than ASM. The method reduces the number of animals needed, reveals important spatial differences in remodelling and could set new analysis standards for murine asthma models.

## Introduction

Asthma is a chronic disease characterised by airway inflammation, hyperresponsiveness and episodes of airway narrowing. Inflammatory events cause the recruitment and activation of eosinophils, mast cells and T-cells, which generate mediators and growth factors amplifying airway inflammation. Bronchoconstrictor stimuli cause acute airway narrowing and additionally promote ASM growth. Repeated episodes of inflammation and bronchoconstriction induce a series of long-term structural changes, comprising increased airway smooth muscle (ASM) mass, extra-cellular matrix (ECM) deposition and epithelial metaplasia; collectively termed airway remodelling, this process results in fixed airway narrowing, increased need for medication and worsening outcomes for people with asthma. Crucially, severe airway remodelling is associated with lung function decline (1).

Physiologic and, more recently, imaging studies have shown that airway remodelling varies with airway size and these differential changes may have important long-term physiologic impacts (2-5). Obtaining tissue samples to study airway remodelling in humans is invasive and generally provides only small tissue samples from large airways at one or two timepoints from which it is difficult to infer whole organ function. Models using repeated airway challenge in sensitised animals are therefore frequently used to understand the mechanisms underlying airway remodelling (6). In the past 10 years, there have been over 900 primary research articles using mice to study airway remodelling which at a conservative estimate equates to 18 000 animals (PubMed search October 2021 -((airway remodelling) AND mouse) AND model AND asthma). Somewhat surprisingly, despite the large number of animals used, there is no consensus on experimental methodology used, nor the type and quality of data reported, and the number with the size of airways analysed are frequently not reported. At best this means that animal tissue is not used to its full advantage and potentially that in some cases a small number of unrepresentative airways are selected, biased by appearance and size. Such methodology also fails to account for, or characterise, intra-subject heterogeneity which is likely to be an important determinant of function at the organ level(7).

Here we set out to develop a methodology which allows us to examine airway remodelling and resolution across the whole range of airways in murine lungs. We hypothesised that this novel approach would reduce selection bias and provide hitherto hidden information on differential structural effects of airway remodelling across airway sizes. Furthermore, by extracting richer data, more information could be obtained from fewer animals and the method may become a standard for both comprehensively assessing airway remodelling and minimising animal use ^(8-12)^.

## Methods

### Animals and Tissue Processing

Ten 6-week old female BALB/C mice underwent ovalbumin (OVA) sensitisation on two occasions 10 days apart. Ten OVA airway challenges were performed between days 17 and 33 as previously described^(13)^. Control animals were sensitised but challenged with phosphate buffered saline (PBS; Sigma Aldrich, UK). Animals were randomised to either the OVA or control challenge groups. OVA-challenged animals were sacrificed by anaesthetic overdose at day 34, 24 hours after the final inhaled challenge, which was defined as maximal remodelling and at days 35, 37, 39 and 41 to track resolution (Fig 1(A)). Control animals were sacrificed at days 34 and 41 only. Group numbers at each timepoint are given in Table 1. For each animal, the trachea was cannulated and bronchoalveolar (BAL) lavage performed using 1ml of chilled sterile PBS. The BAL fluid (BALF) was centrifuged at 1500rpm (4°C) for 10 minutes and the supernatant removed. The pelleted inflammatory cells were resuspended in 1ml sterile PBS, and 200µl of the cell suspension was placed in a Cytofunnel™ (Thermo Fisher, UK) and centrifuged at 450rpm for 6 minutes. The cytospun cells were stained using Rapi-Diff staining kit following manufacturer’s instructions. Slides were visualised using a Nikon Eclipse 90i microscope and the percentage of macrophages, eosinophils, neutrophils and lymphocytes counted by an observer blind to the animal’s treatment.

**Table 1:**
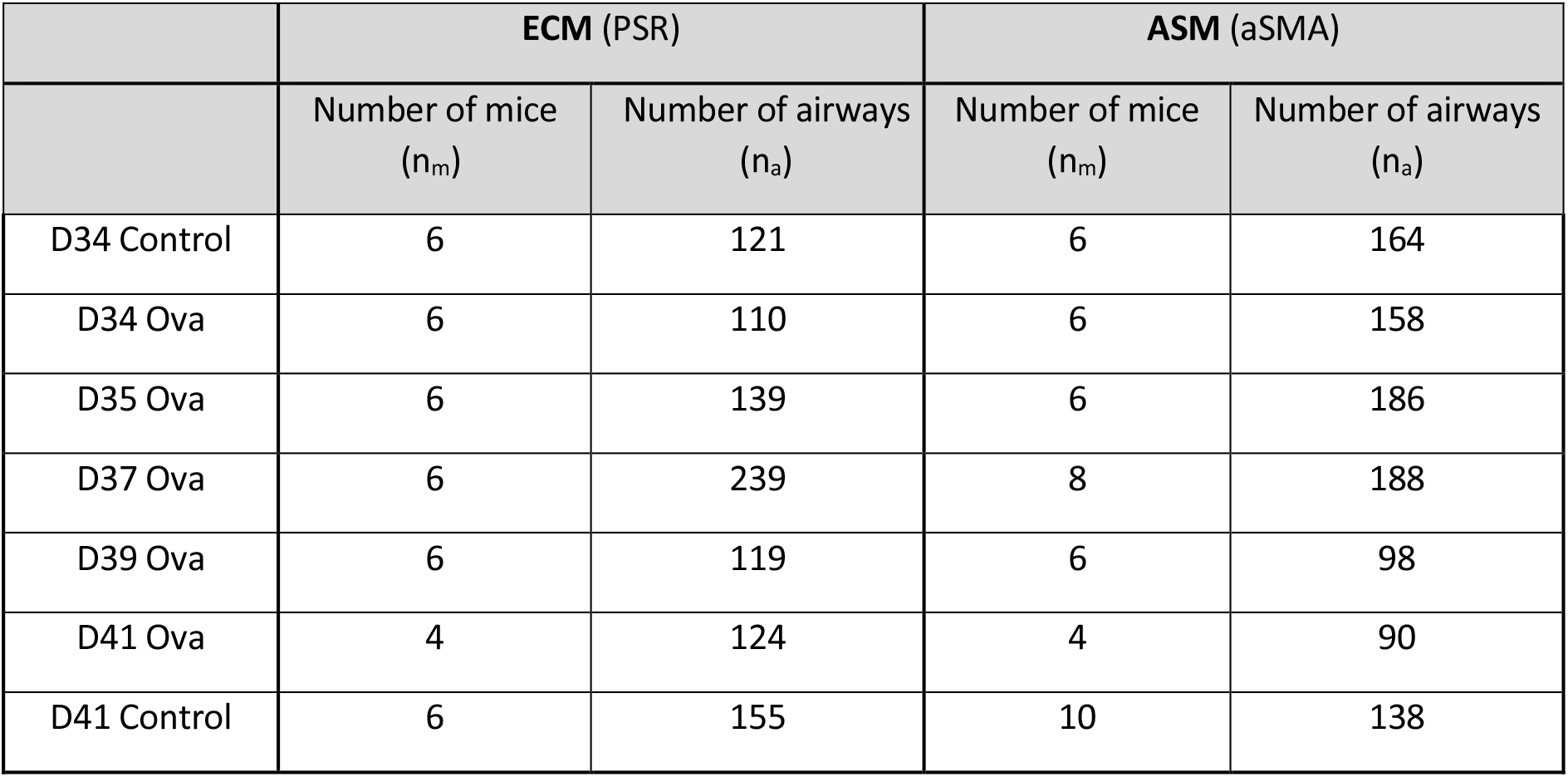
Number of mice and airways identified in each group and at each time point.

**Figure 1:**
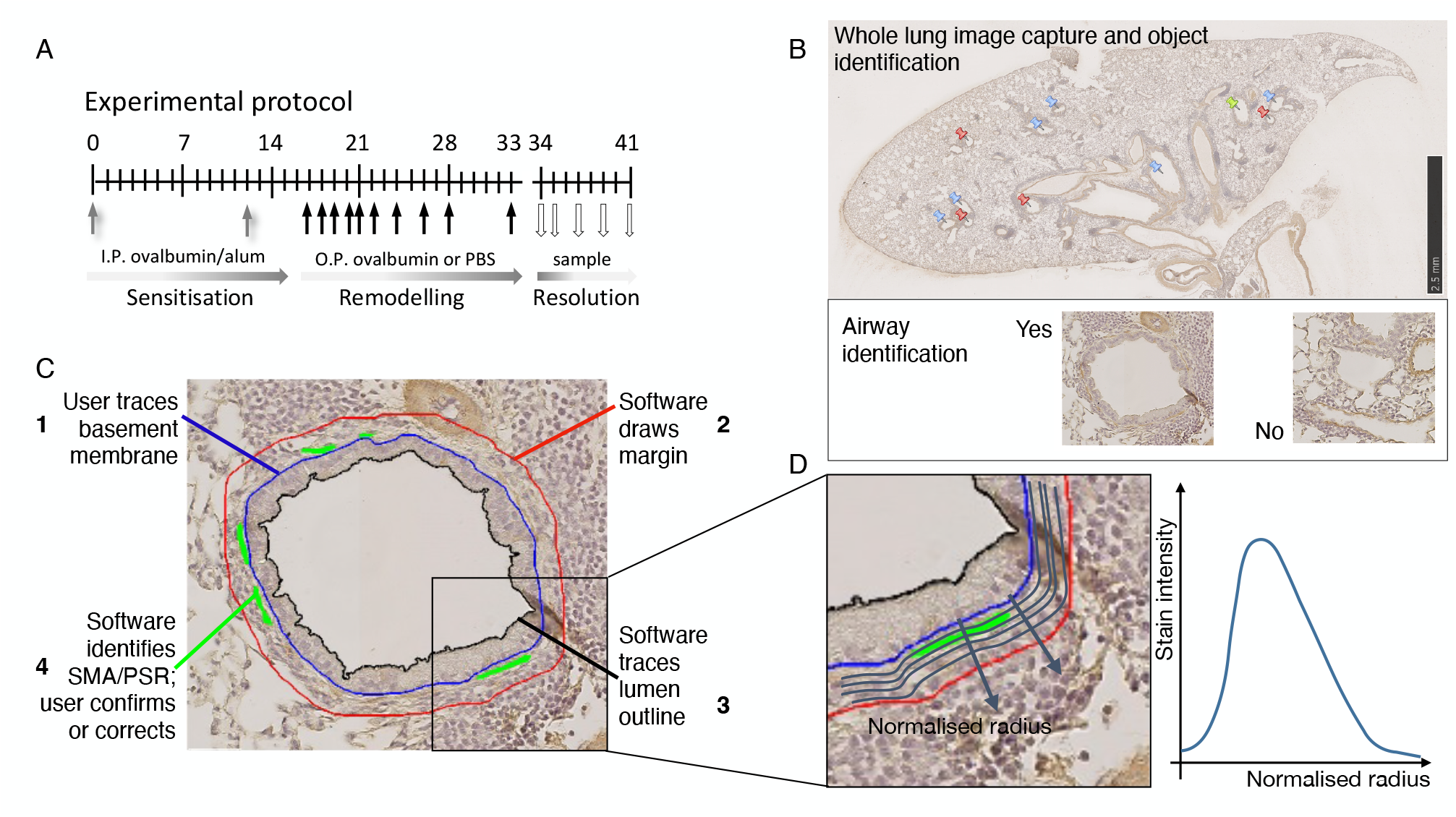
Experimental protocol and data capture. **(A)** Twelve-week old BALB/C mice underwent ovalbumin (OVA) sensitisation on study days 0 and 10 (grey arrows) followed by 10 ovalbumin airway challenges between days 17 and 33 (black arrows) and were sacrificed on day 34 (maximal remodelling) and days 35, 37, 39 and 41 (white arrows). Control animals were sensitised but challenged with physiological buffered saline (PBS) and sacrificed at days 34 and 41. **(B)** Whole lung sections were captured using a Mammatsu Nanozoomer. The software identifies and filters all airway-like objects (indicated with pins) and the user selects the images representing airways from the pool of objects. **(C)** For each airway the user manually traces the basement membrane (BM) (1); the software then draws an outer airway margin (2); the software identifies smooth muscle actin (SMA) or picrosirius red (PSR) stained area which the user confirms or corrects (3) and the software traces the lumen outline (4). **(D)** The spatial distributions of ASM and ECM within the airway are calculated by determining the number of stained pixels within narrow bands between the basement membrane (blue curve) and the band margin (red curve). The number of pixels is averaged over the perimeter of that band and plotted as a function of the normalised radius which places the basement membrane at 0 and the outer airway margin at 1 giving a spatial distribution of stain intensity as shown in the schematic.

The lungs were inflated with 1ml formalin and the whole respiratory tract paraffin wax embedded *en block*. Lung samples were sectioned through their coronal axis and sequential sections stained for α-smooth muscle actin (SMA) by immunohistochemistry and picrosirius red (PSR) as previously described (14) and in supplemental methods. Whole slide images were then captured with a Digital Nanozoomer (Hamamatsu Photonics UK), converted to tiff format at maximum resolution using ndpi2tiff (https://www.imnc.in2p3.fr/pagesperso/deroulers/software/ndpitools/) and imported into MATLAB (The Mathworks Inc) for processing. The custom image analysis software developed in MATLAB is described below (and can be downloaded from: https://github.com/BindiBrook/AirwayIdentification); full details are given in the online supplement.

### MATLAB Workflow and Program Design

A custom pre-processor was developed in MATLAB to identify potential airways which were then filtered to keep objects with characteristics associated with airways and eliminate those without. Full details and code are provided in the on-line supplement and at: https://github.com/BindiBrook/AirwayIdentification respectively. Briefly, irregular circular objects were identified via a multi-step process and converted to a binary image. Epithelial breaks were closed, the lumen identified, smoothed and areas calculated. Minimum and maximum effective diameters were set between 25 and 500 μm, with objects outside the selected range eliminated in order to avoid including cross-sections smaller than alveoli (15) and as large as the main bronchi (16). The perimeter of the trace and the area bounded were computed and objects with pre-defined area-perimeter ratios chosen as potential airways (Fig 1B). A custom filtering algorithm, based upon the airway shape regularity and wall density was used to determine airways from other objects such as blood vessels or alveoli. A script then allowed visualisation of the filtered images for manual review and editing. Objects that were classified as “rejected airways” were also retained for full user checking if required.

### Airway Analysis and Quantification

After airway identification the outer boundary of the epithelium, representing the basement membrane was traced manually for every airway (Fig 1C). The script then automatically placed a second boundary to define the airway wall by dilating the region bounded by the basement membrane by 40 µm. This uniform airway wall thickness was chosen to provide a consistent way to ensure that all airway features are accommodated, without including excessive lung parenchyma, and avoided difficulties associated with identifying the outer boundary of the airway wall in an automatic way. SMA or PSR positivity was quantified by threshold setting of stained pixels within the airway objects. Separate user-defined thresholds were employed for SMA and PSR staining. Total airway area fractions of ASM and ECM was computed as the ratio of the area of SMA or PSR positive pixels respectively, relative to the total area within the basement membrane and second boundary. Spatial distributions across the airway wall thickness were determined similarly, by the ratio of the area of SMA or PSR pixels occurring within a narrow ‘annulus’ between two radial spatial positions, and the total within that region, with these regions being obtained via the dilation process as above (Fig1D). Lumen area and ‘inner area’ (the area contained within the basement membrane outline) are determined by pixel count, as above; epithelial area was determined by subtracting the lumen area from the inner area in the SMA-stained airway images (Supp Fig 6A).

### Data Analysis and Statistics

Data analysis and statistics were performed using custom scripts in MATLAB. The complete datasets are available <u>here</u>.

### Study approval

Animal work was approved by the Animal Welfare and Ethical Review Board (AWERB) of the University of Nottingham (UK) and conducted in accordance with all terms of the Establishment, Project and Personal Licenses issued by the Secretary of State for the Home Office. Consistent with all national and international law, studies were carried out as detailed in the Animal [Scientific Procedures] Act 1986 (Amended Regulations 2012) (ASPA), Animal Welfare Act 2006, Directive 2010/63/EU, the LASA guidelines and in respect to the principals of Replacement, Reduction and Refinement. Work was performed under license number RGJ 40/3709 using the 19b5 protocol.

## Results

At the end of the 34-day sensitisation and challenge protocol, in the saline challenged control mice macrophages comprised 99% (SEM±0.5) of BAL cells. Chronic OVA challenge was associated with an increase in BAL eosinophils (43±3.5%), lymphocytes (13±1.6%), neutrophils (4±0.9%) and a relative reduction in macrophages (40%±3.6). These changes progressively and completely resolved over seven days following the cessation of airway OVA-challenge (Figure 2A).

**Figure 2:**
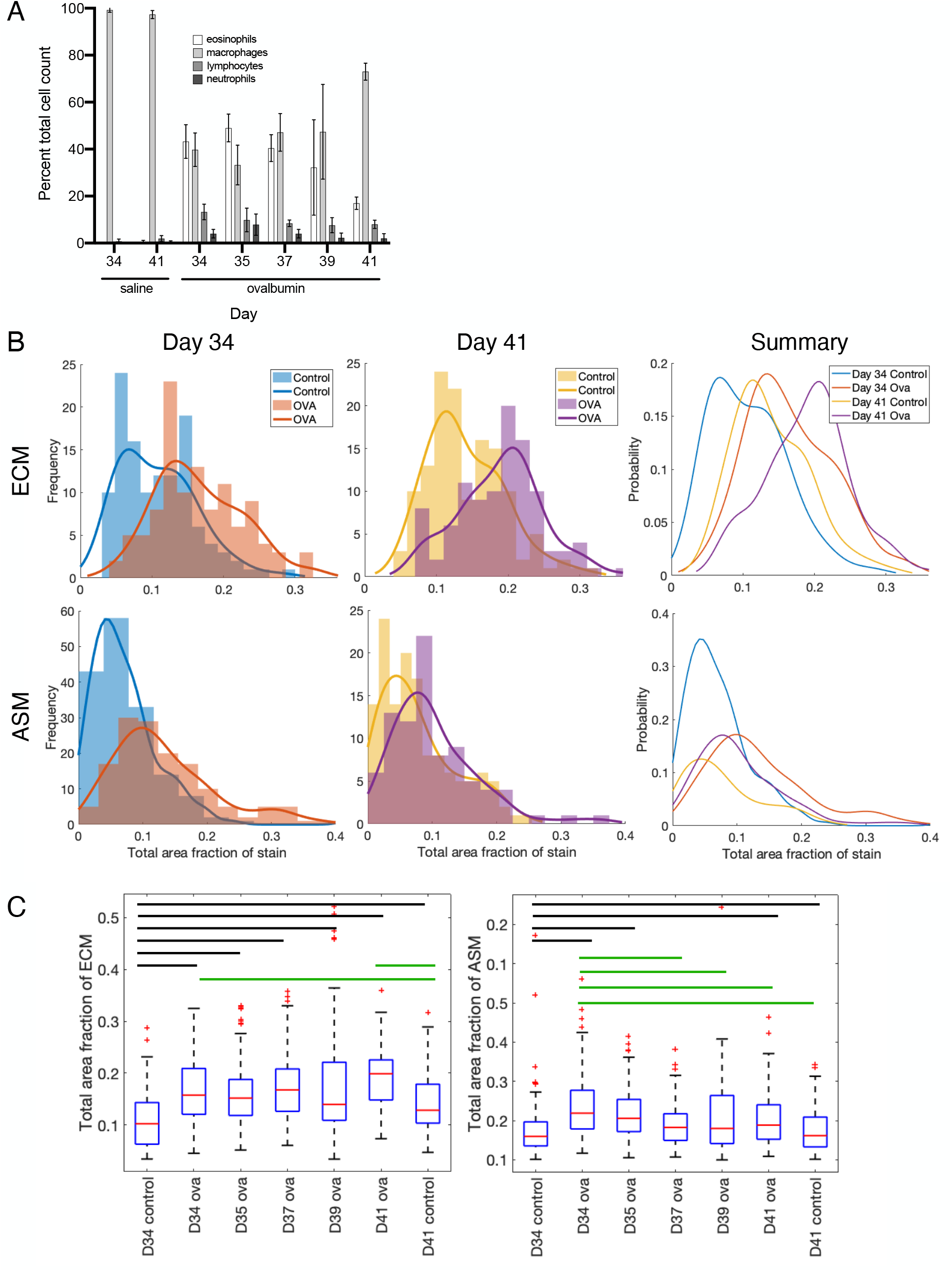
Global changes in smooth muscle and extra-cellular matrix during remodelling and resolution. **(A)** Bronchoalveolar lavage cell count. Eosinophils, macrophages, lymphocytes and neutrophils as a percent of total number of cells for both control and ova-challenged animals on days 34 and 41 and for ova-challenged animals on days 35, 37 and 39. **(B)** Area fractions of extra-cellular matrix (ECM; top row) and airway smooth muscle (ASM; bottom row) for all airways from both control and ova-challenged animals at maximal remodelling (day 34) and one week post challenge (day 41). Histograms and corresponding fitted distributions of area fraction of stain at days 34 (left column), and 41 (middle column). All four fitted distributions are shown in the right column (data from 4-6 mice and 90-168 airways for each condition – see **Table 1** for group numbers). **(C)** Global changes in airway constituents during resolution. Box plots show change in area fraction of ECM (left) and ASM (right) between days 34 and 41. Box shows the median, 25^th^, 75th percentiles and whiskers the most extreme data points not considered outliers. Black bars indicate means significantly different to day 34 control. Green bars indicate means of ECM fraction that are significantly different to day 41 control (left) and means of ASM area fraction that are significantly different compared with mean of day 34 ova (right). Analysed using one-way ANOVA; p= 2.5e-23 and p = 5e-15 respectively.

### Global changes in airway smooth muscle and collagen

The protocol identified a total of 1007 PSR-stained and 1022 SMA-stained airways across all 28 control and 58 OVA-challenged animals over 5 timepoints. At each timepoint we identified 90-240 airways (denoted *n*_*a*_) from 4-10 mice (denoted *n*_*m*_) (see Table 1 for breakdown). ECM and ASM area fractions were calculated for each airway (Supplementary Table 1). To determine if differences between individual animals and group sizes affected the data interpretation, we calculated: (i) the mean ECM and ASM area fraction per mouse (denoted μ_m_), (ii) the group mean ECM and ASM area fractions for all mice across all the *n*_*a*_ airways at each timepoint (μ_a_), and (iii) the mean ECM and ASM area fractions for each group of animals (μ^*^_m_) obtained by taking the average over the *n*_*m*_ values, μ_m_. We found that μ_m_ and μ^*^_m_were very similar, and did not differ significantly from the group mean μ_a_. Therefore, most of the data are presented using grouped airways (i.e., *n*_*a*_, μ_*a*_) rather than individual animals.

Using all 553 airways identified at the point of maximal remodelling on day 34, we initially tested the effect of examining the data as grouped airways, rather than collapsing all values to a single mean per mouse. The area fractions of ECM and ASM for OVA-challenged and control animals were first calculated as group means (μ_*a*_), where we observed a 1.5 and 1.8 fold increase in the mean area fractions of ECM and ASM in OVA-challenged, relative to control animals (one-way ANOVA p = 2.5×10^−15^, 5×10^−15^ respectively; Fig 2C). These differences in ASM and ECM area fractions were confirmed when performing the same analysis with means obtained from individual animals (μ_*m*_* Supp Table 1). Therefore, by considering airways individually (*n*_*a*_) we can retain the rich information on intra-subject heterogeneity which would otherwise be lost (Supp Fig 4). Next, we considered the distributions of these area fractions within each group (Fig 2B). On day 34, ASM and ECM area fractions increased in OVA-challenged when compared with control animals (as shown by the right shift in distribution peak. (Figure 1). Additionally, the heterogeneity of ECM and ASM remodelling (as highlighted by the width of the distributions) differed significantly from control airways with a 1.3 and 1.5-fold increase in variance respectively (Fig 2B).

### Global changes in airway constituents during resolution period

We next examined the resolution of these changes over time in the 776 PSR and 700 SMA-stained airways identified at days 34 and 41 in control and OVA-challenged animals. The mean ASM area fraction in OVA-challenged animals, observed at day 34, was significantly greater than in control animals (one-way ANOVA p=5×10^−15^). Over seven days from the last airway challenge on day 34, the ASM area fraction decreased progressively to almost complete resolution at day 41 (Figs 2B and C). The variability in ASM area fractions associated with remodelling also decreased to similar levels to control animals by day 41 in all but those airways with the highest ASM fractions (Fig 2B).

The mean ECM area fraction in OVA-challenged animals, observed at day 34, was significantly greater than in control animals (one-way ANOVA p=2.5 × 10-23) but, in contrast to ASM, remained elevated over the seven days after cessation of airway challenge. The increased variability in the ECM area fractions associated with remodelling was also sustained (Fig 2C). Additionally, ECM fraction increased 1.3-fold in control animals between day 34 and 41 although these changes were smaller than those due to OVA challenge-associated remodelling.

### Stratification of remodelling changes by airway size

The methodology allows detailed measurement of additional airway parameters including lumen area and basement membrane (BM) perimeter. This, and the large number of airways studied, allows examination of the pattern and time course of remodelling and resolution across airways of different sizes for the first time. Airways were initially arbitrarily divided into small (BM perimeter <500μm), medium (501μm-1000μm) and large (1001μm-1500μm) categories. To allow us to make direct comparisons between airways stratified by size, we first determined that both the mean BM perimeter and their range of distribution at day 34 did not differ between control and OVA-challenged animals (one-way ANOVA, p= 0.166 and two-sample KS tests, p=0.08 respectively). At day 41 mean BM perimeter did not differ significantly (one-way ANOVA, p=0.166), although the distribution of these changes did vary somewhat (two-sample KS tests p=0.0069).

In OVA-challenged animals at maximal remodelling on day 34, mean ECM area fraction increased significantly in small and medium sized airways by 1.7 and 1.4-fold respectively compared with control animals. A 1.3-fold increase in large airways was not significant (Fig 3A). ECM remodelling in each category was sustained over the post-challenge period to day 41 with 1.9, 1.5 and 1.4-fold increases respectively, compared with control at day 34 (Fig 3A). Mean ASM area fraction also increased at day 34 in all airway sizes, although the pattern of remodelling differed with the greatest increase in large airways; 1.8, 1.5 and 1.9-fold increases were observed in small, medium and large airways, respectively (Fig 3B). The mean ASM area fraction at day 41 had fallen to control levels in medium and large but not small airways; however, only a relatively small number of airways could be identified in this group (Fig 3B). ASM remodelling resolved to a greater degree in large and small rather than medium-sized airways (Fig 3B, middle column) and is consistent with the wider distribution of ASM fractions at day 41 in the OVA-challenged animals (Fig 2B).

**Figure 3:**
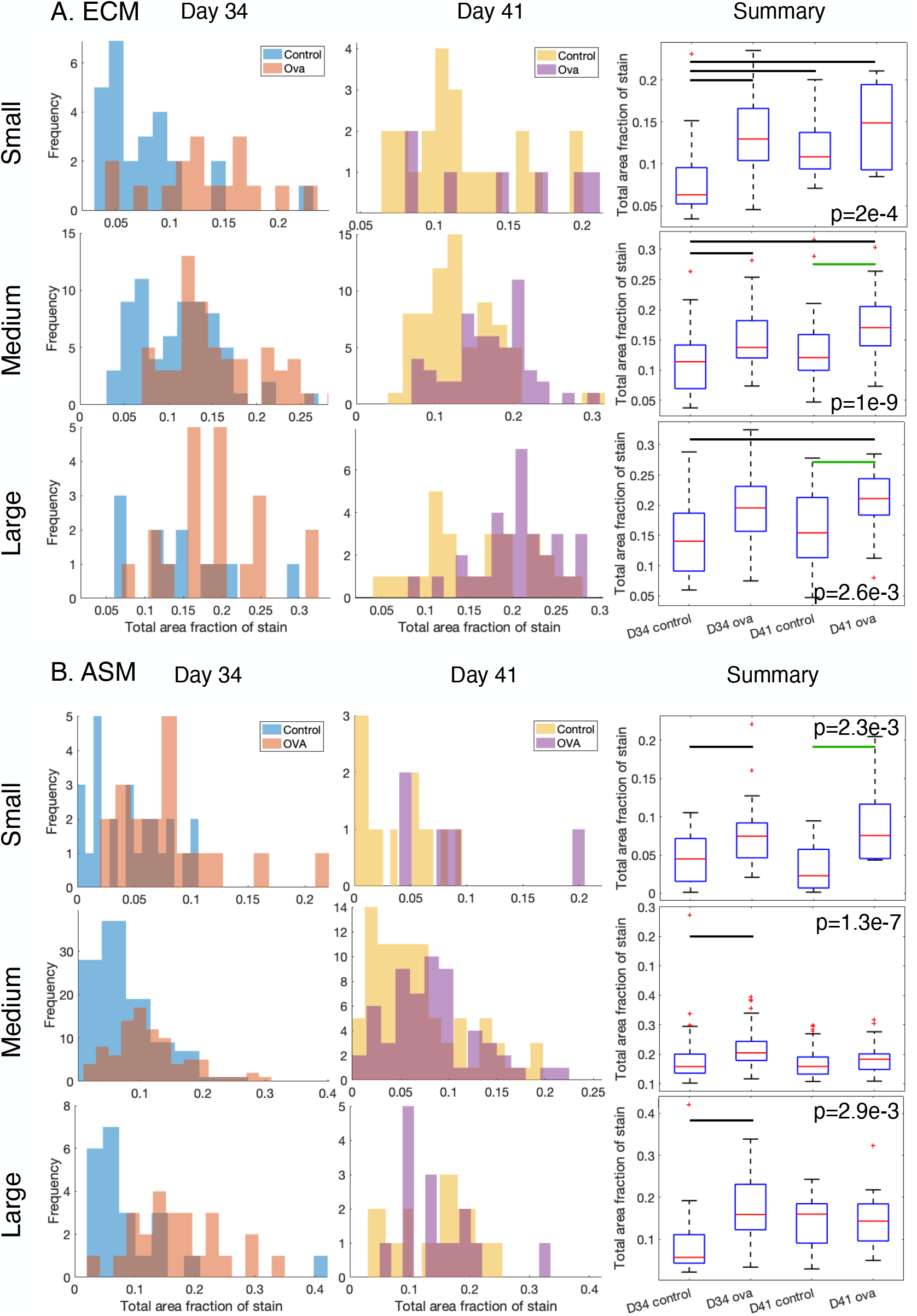
Changes in smooth muscle and extra-cellular matrix stratified by airway size. Airways were categorised as small (basement membrane perimeter < 500µm), medium (500 – 1000µm) and large (1000 – 1500µm); numbers of airways identified in each these groups are given in **Table 2**. Histograms show area fraction of **(A)** extracellular matrix (ECM) and **(B)** airway smooth muscle (ASM) at days 34 (left column), and 41 (middle column). Box plots show change in area fractions of ECM and ASM (right column) on days 34 and 41. Box shows the median, 25^th^, 75th percentiles and whiskers the most extreme data points not considered outliers. Black bars indicate means are significantly different to mean at day 34 control. Green bars show cases in which there are additional significant differences between means. Analysed using using one-way ANOVA; p values shown on figure panels.

**Table 2:**
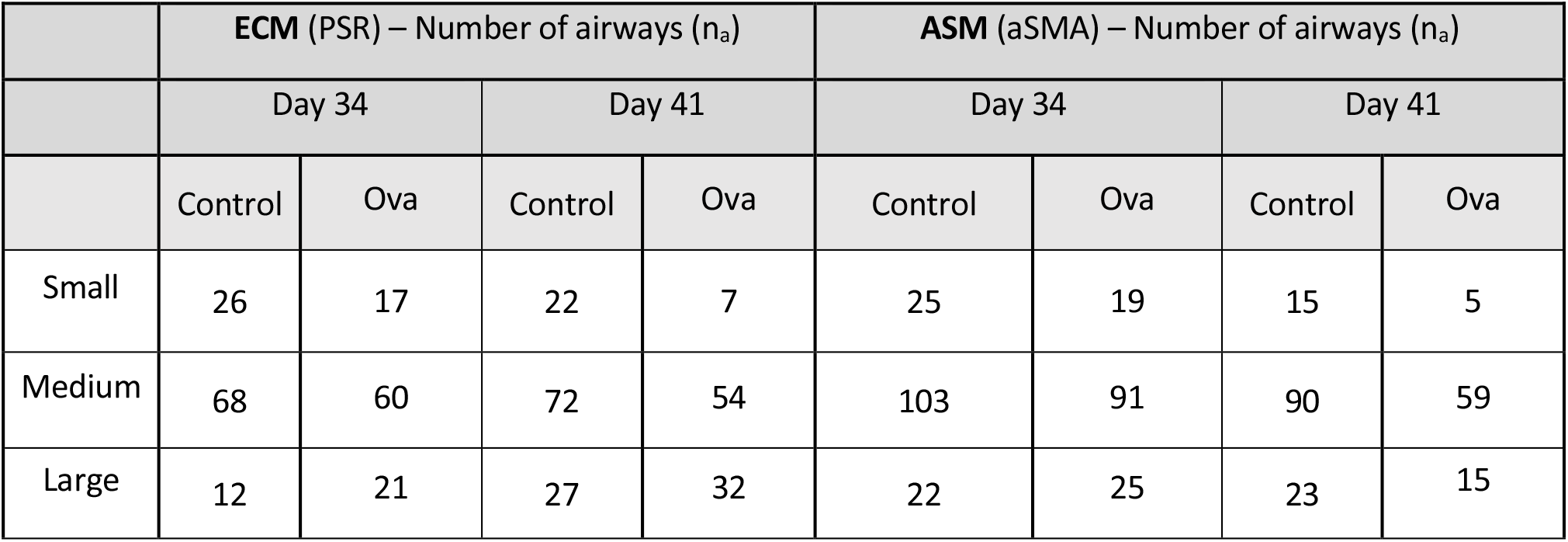
Number of airways identified in each group stratified by airway size (small (basement membrane perimeter < 500µm), medium (500 – 1000µm) and large (1000 – 1500µm)) at Days 34 and 41.

### Fine stratification by airway size

These airway size categories are a convenient way to observe differential airway remodelling and highlight how the distributions within each airway range informs their interpretation. However, our dataset allows us to examine the heterogeneity in airway remodelling in finer detail by considering the relationship between ASM and ECM fractions across the whole range of airway sizes (defined by basement membrane perimeter). As BM perimeter distributions do not differ between control and OVA-challenged animals at day 34, the remodelling observed can largely be attributed to increased airway ASM and ECM content. 3D histograms showing the frequency of airway area fraction split across finely-resolved size categories, show that ECM and ASM distributions are more diffuse in OVA-challenged compared with control animals, consistent with the increased variance seen with airway size stratification (Fig 4). The changes in orientation of the distribution peaks, shown by lines of best-fit (Fig 4 and Supplementary Figure 5) highlight the airway size-dependence of remodelling in more granular detail. Consistent with our findings based on coarse stratification, we observe that ECM remodelling decreases progressively with increasing airway size, with little to no remodelling observed in airways with BM perimeter greater than 1000µm. For ASM distributions, however, there is a significant change in peak orientation, corresponding to increased remodelling in small compared with larger airways (with BM greater than approximately 750µm), although the small number of large airways means that the linear correlation lines should be interpreted with care in the latter region. The resolution of remodelling in both ECM and ASM, shows substantial variation over the seven-day post-challenge period highlighted by tracking the location of mean ECM and ASM from the 2D projections over all time points (Supp Fig 5E-H). This emphasises the contrast in the overall trend for ASM content to resolve to almost control levels, while ECM continues to remodel throughout this period.

**Figure 4:**
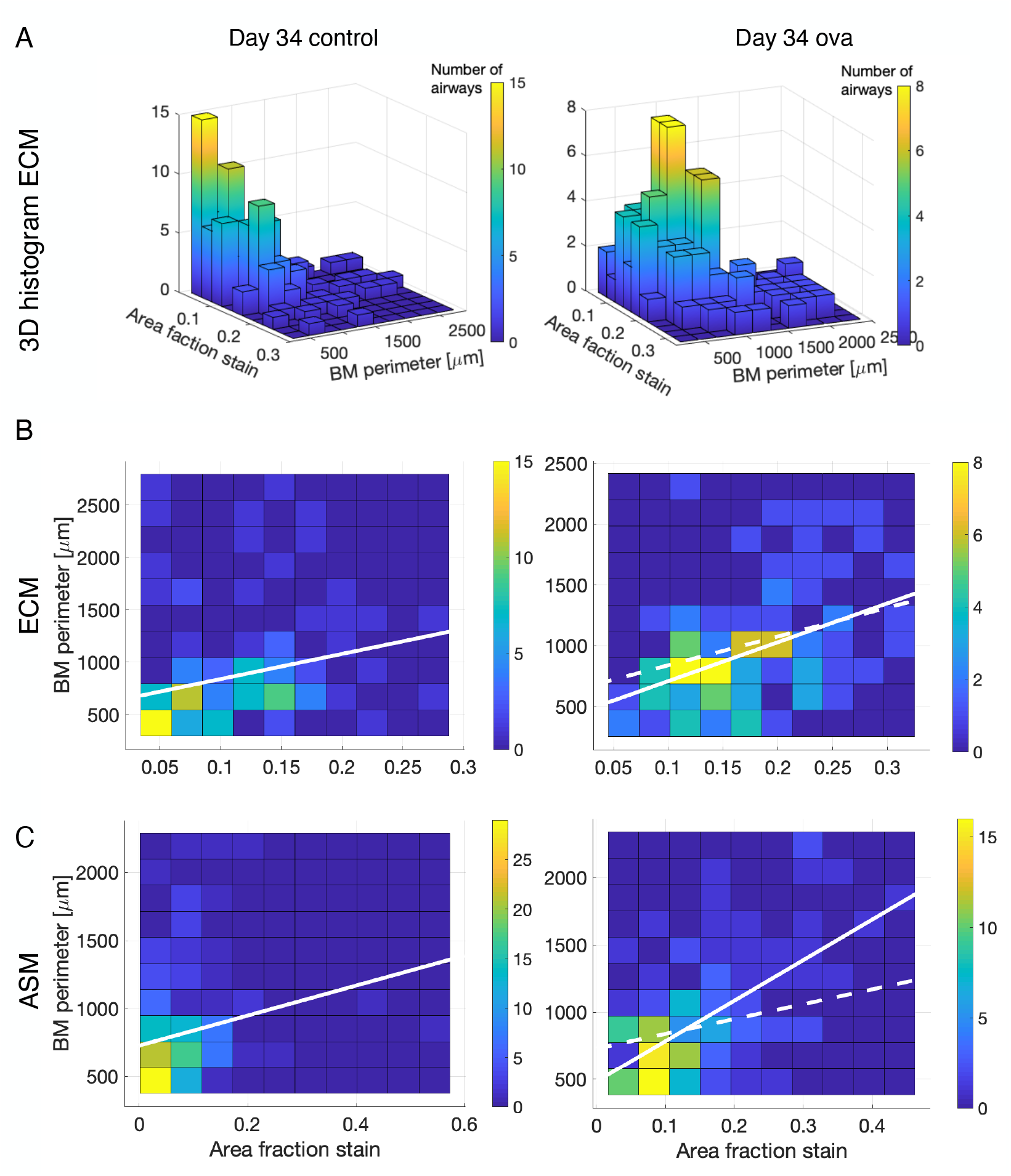
Differential changes in airway constituents with airway size under fine stratification. **(A)** 3D histograms showing frequency of airways (denoted by colour-scale) binned according to both ECM area fraction and size (basement membrane (BM) perimeter). To aid interpretation 2D projections (top-down views of the 3D histograms) are shown instead for **(B)** Extracellular matrix (ECM) and **(C)** airway smooth muscle (ASM) for day 34 control (left column) and day 34 ovalbumin challenged (ova, right column). Solid white lines indicate lines of “best fit”. Dashed white lines are lines of best fit from control data overlaid on the corresponding ovalbumin data for comparison. Interpretation of these, and data for all days are given in **Supplementary Fig. 5**.

### Spatial distributions of ASM and ECM across the airway wall

We next examined how total area fraction of ASM and ECM were distributed radially from the basement membrane to the outer airway margin (Fig 1D), and how this changed during remodelling (Fig 5). In control animals at day 34 and 41, ECM content was highest in the 40% of the airway adjacent to the basement membrane, peaking at 20% in each case (blue and yellow curves, Fig 5A, left column; Fig 5C, left panel). In contrast, increased ECM content in control animals at day 41 was restricted to the 30% of the airway wall adjacent to the basement membrane, with little or no increase elsewhere. ECM remodelling in OVA-challenged animals at day 34 followed a similar pattern, but with a shift in the peak location towards the airway wall mid-point (Fig 5A, top right panel) and increased ECM deposition in the outer 80% of the airway compared with control at day 34 (cf. blue and orange curves in Fig 5A (top row) and Fig 5C (left panel)). The increased ECM observed in outer regions of the airways in OVA-challenged animals at day 34 is maintained during the post-challenge phase until day 41 (cf. orange and purple curves in Fig 5A and Fig 5C (left panel)). Further increases in ECM content occurred in the 40% of the airway adjacent to the basement membrane and, interestingly, the location of the peak in the OVA-treated animals shifted back towards the basement membrane, as exhibited by the control data (cf. purple and yellow curves in Fig 5C, left panel). ASM content in control animals was also highest in the 40% of the airway thickness nearest the basement membrane, peaking at 20% with very little ASM more peripherally (Fig 5B, left column). OVA-challenge resulted in increased ASM content in the region 20-80% from the basement membrane, with a shift in the peak further away from the basement membrane (cf. blue and orange curves in Fig 5B (top row) and Fig 5C (right panel)). Over the post-challenge period the peak and overall distribution, returned close to control levels (cf. purple and yellow/blue curves; Fig 5C (right panel)).

**Figure 5:**
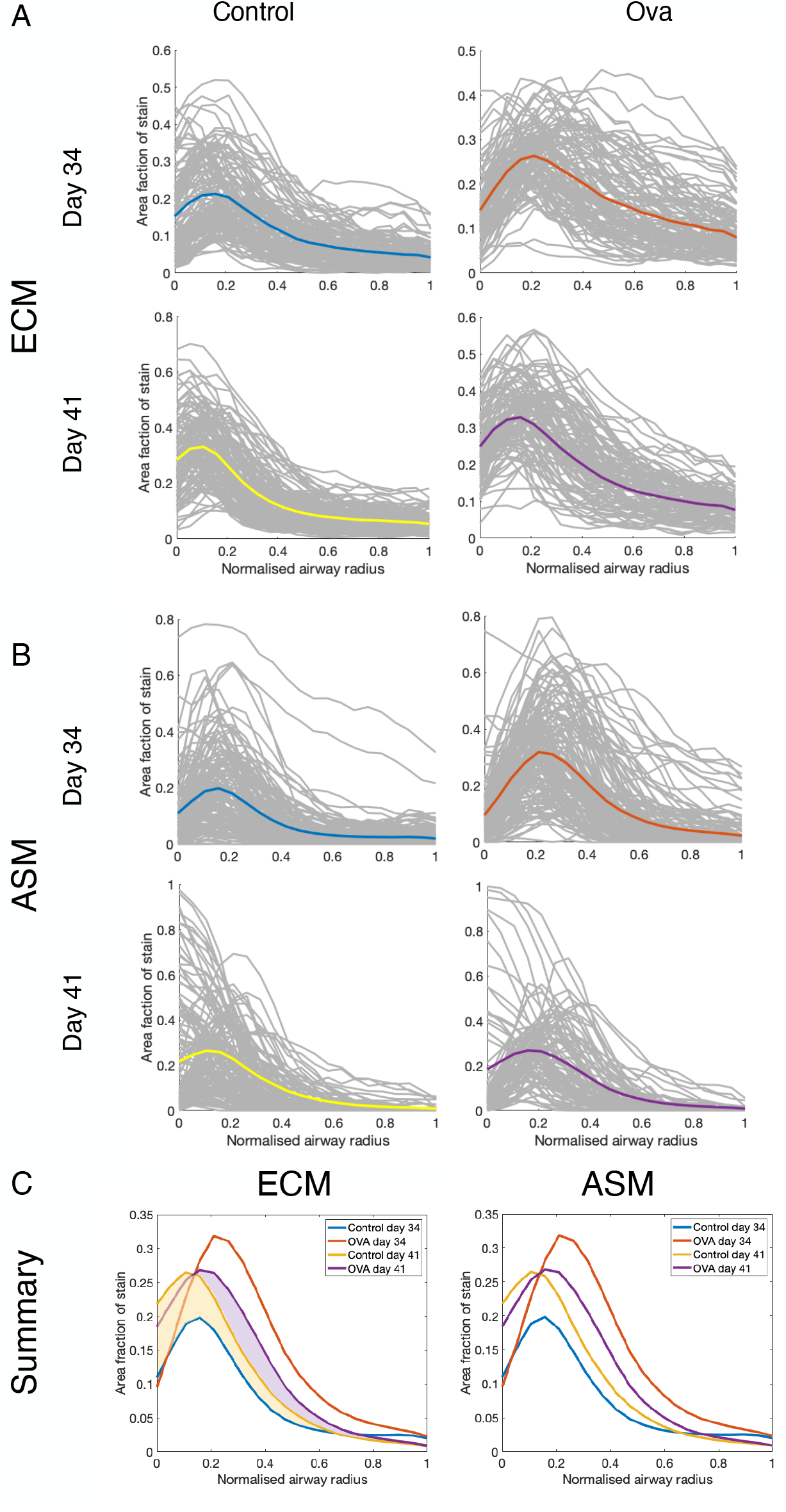
Spatial distribution of smooth muscle and extra-cellular matrix across the airway wall. Area fraction of **(A)** extra-cellular matrix (ECM) and **(B)** airway smooth muscle (ASM) distributed spatially across the airway wall for all control and ovalbumin-challenged (Ova) mice at days 34 and 41. The region between the basement membrane and the outer airway margin defined by the software is scaled from zero (basement membrane side) to one (outer airway margin). Gray curves represent individual airways, and coloured curves indicate average area fraction as function of normalized airway radius and shown in summary in **(C)**. The yellow shaded region indicates increased ECM between control d34 and control d41 (not significant). Purple shaded region shows increase in ECM content between ova d41 and control d41 (significant). Distributions for individual mice are given in **Supplementary Fig. 4**.

The comprehensive dataset, shown in Figs 4 and 5A&B, also highlights the heterogeneity in airway composition both in control and post-challenge conditions. We further investigate this in Supplemental Figure 4, demonstrating wide variation in total ECM area fraction and its radial variation between animals, but with somewhat more consistent response observed in OVA-challenge conditions compared to control. Additionally, BM perimeter shows greater consistency between animals than does airway constituent remodelling.

### Epithelial involvement in remodelling

To determine how the epithelial layer was affected by remodelling (Fig 6) we computed the epithelial area for each airway as shown in Supplementary Figure 6A. To correct for airway size, the epithelial area between the BM and airway lumen was normalised to the BM perimeter. For 550 airways examined at days 34 and 41, the normalised mean epithelial area at day 34 was 1.4 fold higher in ovalbumin-challenged compared with control animals (p = 1.2×10^−32^). In the 472 airways studied at days 35, 37 and 39, the mean epithelial area was also significantly increased compared with control at day 34 (Fig 6B). Epithelial area remained elevated at day 41 with a 1.4-fold and 1.5-fold increase over day 34 and 41 control animals respectively (Fig 6A and B). In addition, variability in epithelial area was increased in the OVA-treated animals. Stratification by size shows that the changes at day 34 were driven by large (1.7-fold; p = 2.4e-8) and medium sized (1.3-fold; p=1.2×10^−19^) but not small airways.

**Figure 6:**
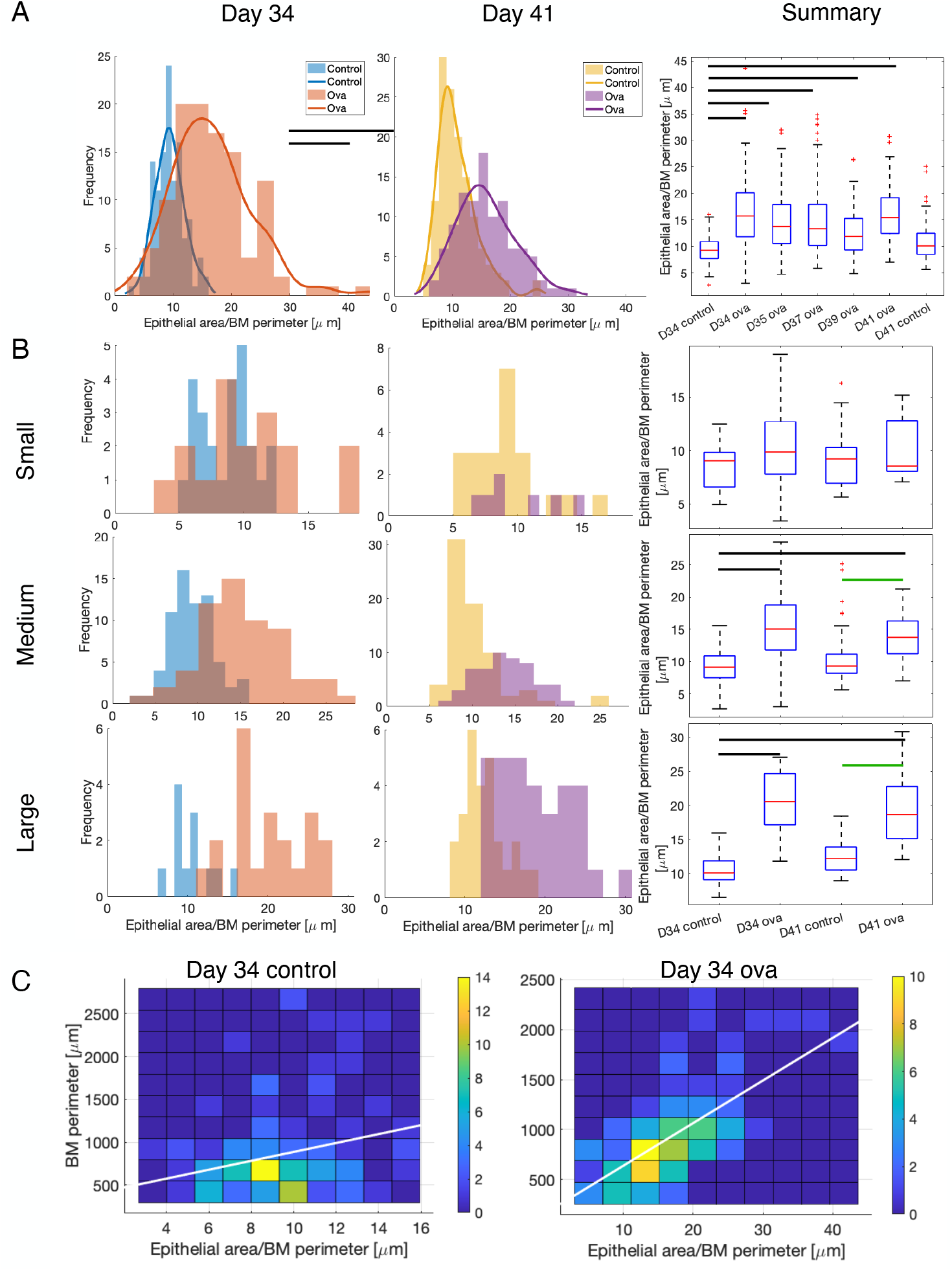
(A) Changes in epithelial area at days 34 and 41. Epithelial area normalized with respect to basement membrane (BM) perimeter for all airways from both control and ovalbumin-challenged (ova) animals. Histograms and corresponding fitted distributions of epithelial area at days 34 and 41 **(B)** Global changes in epithelial area during resolution period. Box plots showing change in epithelial area from days 34 to 41. Box shows the median, 25^th^, 75th percentiles and whiskers the most extreme data points not considered outliers. Black bars indicate means that are significantly different to mean at day 34 control (p values indicated in text; 4-6 mice and 90-186 airways per group -see **Table 1** (aSMA) for group numbers). **(C)** Changes in epithelial area graded by airway size. Airways were categorised as small (BM perimeter < 500µm), medium (500 – 1000µm), and large (1000 – 1500µm). Histograms show epithelial area at days 34 and 41 (see **Table 2** (aSMA) for group numbers). **(D)** Summary data box plots showing change in epithelial area on days 34 and 41 corresponding to the airway size. Black bars indicate means are significantly different to control at day 34. Green bars show additional significant differences between means. (E) Differential changes in epithelial area with airway size under finer stratification. 2D projections of 3D histograms showing frequency of airways (denoted by colour-scale) binned according to both epithelial area and size (basement membrane perimeter) at day 34. White lines indicate lines of “best fit”. Interpretation of these are given in **Supplementary Fig. 5**.

We expected the observed increase in epithelial area to result in a decrease in normalised lumen area, given that the mean BM perimeter remains largely unchanged. However, the mean lumen area was not significantly altered in OVA-challenged compared to control animals at either day 34 or 41 (one-way ANOVA, p=0.9935, 0.0616 respectively). This observation was investigated further using simulations based on day 34 control data (Fig 7), together with simplified geometrical arguments (Supplementary figure 6). The latter provides a mathematical constraint under which, both epithelial and lumen area in OVA-challenged animals can indeed exceed epithelial and lumen area in control animals. The former employs a simple statistical model under which the lumen and inner (that enclosed by the basement membrane) area data in OVA conditions is obtained by applying small random perturbations to control data, with epithelial area being computed from their difference. Full details are given in the Supplementary material. Simulated distributions have similar shapes to that of the observed experimental data in OVA-challenged animals at day 34 (cf. green and orange curves in Figs 7A&B, left column; note that the distribution amplitude is not expected to be similar, since these data are directly computed from the control (blue curves)), with similar significant and non-significant increase in means (Figs 7A&B, right column). The corresponding distribution of simulated epithelial area also shows a significant increase in mean and a similar shape to the observed data in OVA-challenged animals at day 34 (Fig 7C). Taken together, these data support and explain the unexpected relationship between epithelial area, BM perimeter and lumen area observed in the data.

**Figure 7:**
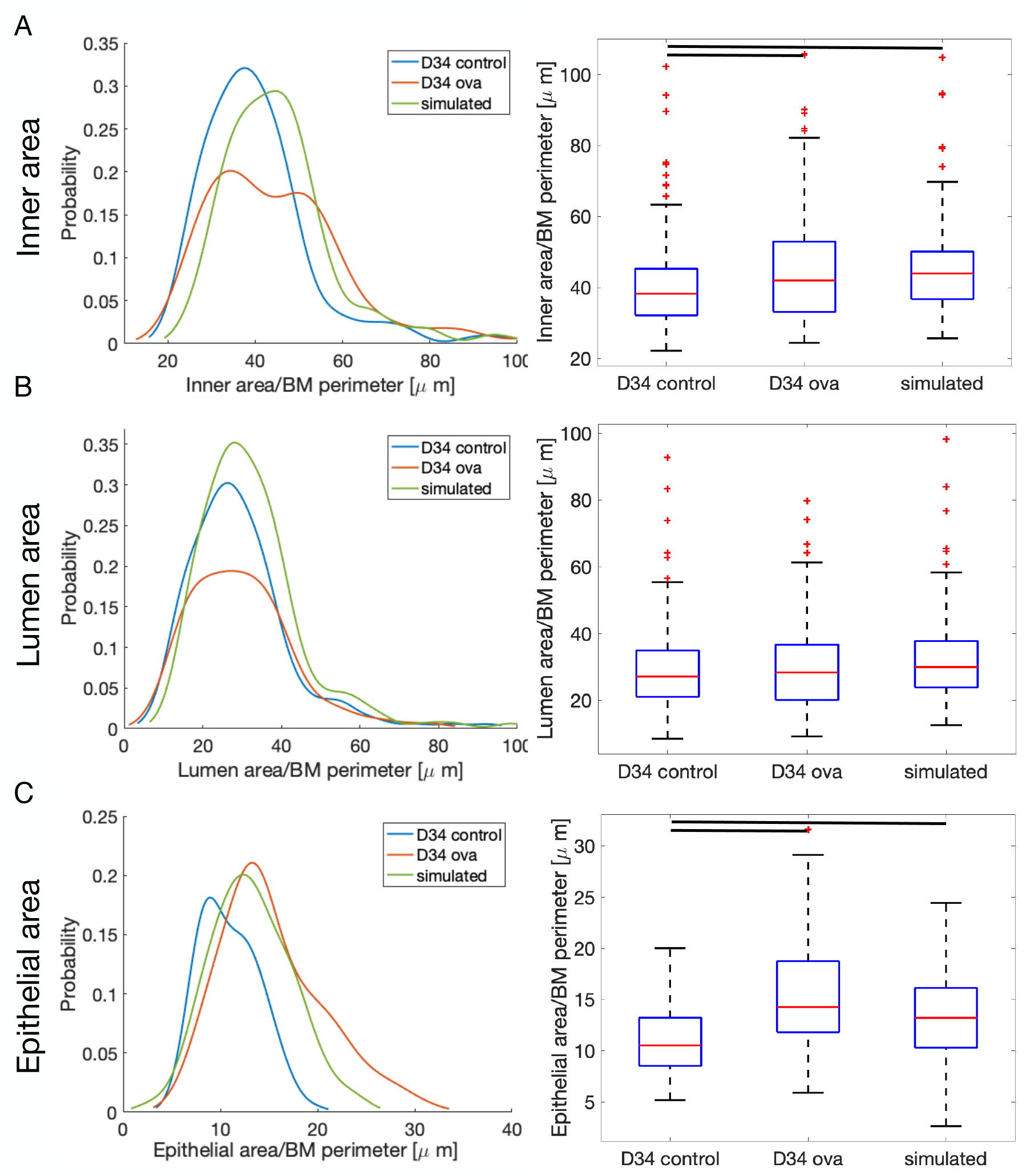
Airway area relationships explained by simulated data. Inner, lumen and epithelial areas are computed as described in Supplementary Figure 6. Simulated lumen and inner area OVA data is obtained by perturbing control data at day 34 by a normally-distributed random amount (see Supplement). Panels show area distributions normalised with respect to the BM perimeter for control and OVA data and simulated OVA data at day 34 (left), and corresponding box plots (right). (A) Inner area normalised with respect to the BM perimeter is significantly increased in ova compared with control at day 34, indicated by the shift in the peak in the distributions (left) as well as one-way ANOVA and box plots (right). (B) Lumen area is not significantly different. (C) Mean epithelial area is significantly increased. Analysed using using one-way ANOVA; p values given in text.

## Discussion

We have developed a new method to examine airway remodelling in mouse whole lung cross-sections. In conjunction with immunohistochemistry, the method allows assessment of the amount and distribution of airway components and can be stratified quantitatively according to airway size. In this study we demonstrate that the method is effective in examining ASM, ECM and epithelial remodelling and for the first time has provided evidence of differential changes in large and small airways in response to chronic airway challenge in mice. Our new technique represents a step-change in airway remodelling analysis over conventional methodology. Exploiting image processing techniques, our method permits semi-automatic identification, and detailed quantitative analysis, of significantly larger numbers of airways from the same number of experimental animals than has hitherto been possible. These extensive datasets support unique insights into airway remodelling processes.

It has long been suggested that remodelling of airways in asthma is not homogeneous and that differential remodelling occurs across airways of varying sizes (17). The implications of heterogeneous remodelling across larger, proximal and more distal small airways are not fully understood. It has been suggested that remodelling of small airways contributes to airway hyper-responsiveness (18), which is consistent with regional changes in ventilation suggestive of distal airway remodelling (19). Supporting this, small, airways significantly contribute to increased total lung resistance in moderate-severe asthma (20). Changes in the distribution and extent of remodelling across airways of varying sizes are also seen between non-fatal and fatal asthma (21) suggesting these observations are clinically important.

The method described here identified profound changes in ASM, ECM and epithelial cell remodelling between airways of varying sizes and demonstrated that distinct features of remodelling do not change in a universal way. Our findings show that small airway remodelling is primarily associated with increases in ASM and ECM fractions whereas epithelial cell remodelling occurs primarily in the larger airways. Increased ASM and ECM mass around small airways in human asthma and animal models has been previously well documented (13, 22-27). However, none of these previously published studies have directly compared ASM and ECM remodelling around small versus large airways. Studies assessing epithelial remodelling in large and small airways are more limited in number; however, Carroll et al. found no significant difference in epithelial damage between large and small airways in human asthma (21). Our study also shows that the resolution of ASM over the 7-day timescale appears to mirror the resolution of eosinophils which occurs over the same timescale. This has previously not been demonstrated; the only other study that we are aware of in which resolution of inflammatory cells was investigated after cessation of challenge (28) did not include time-points as finely resolved as in our study.

A strength of the method presented here is the ability to assess features of remodelling across airways of all sizes in an unbiased manner, which has not previously been possible in other studies of airway remodelling. Airway remodelling is primarily assessed using histological and immunohistochemical approaches, which while being excellent methods to study pathological changes in tissues, are prone to many types of selection bias for a number of reasons (29). The ability to scan an entire transverse section of lung tissue, identify every airway present and make multiple measurements within those airways reduces selection and sampling bias. A previous attempt to measure remodelling in airways of all sizes within lung tissue was limited by the dependence on the experimenter to identify airways and draw their boundaries reducing the precision in defining airway size (22). Our system represents a significant advancement in the automatic identification of all airways present and accurate calculation of airway wall dimensions once the basement membrane has been defined manually. This is, to our knowledge, the first automated system for identifying airways within lung tissue. This model has the potential to set a new, higher standard for the analysis and reporting of airway remodelling changes in animal models of asthma.

A second strength of the method presented here is the ability to assess spatial changes in ASM and ECM occurring within the airway. Spatial distributions suggest that although ECM close to the membrane increased in control animals at day 41 compared with control animals at day 34, the more diffuse increase in ECM in the rest of the airway occurs only in OVA-challenged animals and not in control animals. This finding suggests that there is some natural ECM increase near the basement membrane through natural ageing of the animals (but that increase is not significant), while the OVA-driven increase is larger and more diffuse. Additionally, we observe that in control animals, areas of higher ECM density appear to correlate with areas of higher ASM density whereas in OVA-challenged animals, increased ECM appears at the outer margins. Taken together, these observations suggest a possible mechanism that warrants further investigation: during airway remodelling increased ASM deposits ECM throughout the airway which then resolves but leaves the ECM behind.

While reporting data for individual animals is largely seen as the accepted standard, we observe little difference in changes when grouping airways for individual animals and assessing airways individually. Our data show that by examining all airways independently, detailed data on intrasubject heterogeneity is preserved that is lost if data is averaged for individual animals. Additionally, the number of airways captured dramatically increases statistical power reducing the number of animals required to appropriately test a hypothesis. Here we have assessed ASM, ECM and epithelial cell remodelling. However, the method allows for a diverse range of measurements to be made, only limited by the availability of a specific immunohistochemical stain, making this a powerful tool to study airway biology and disease.

While the method has many advantages there are some limitations. At present the automatic airway identification requires user confirmation that the object identified is an airway, which is somewhat time-consuming. Future work by this group will explore the use of artificial intelligence and machine learning to classify airways and segment stained tissue, removing the need for user-dependent airway identification. Finally, due to the nature of the airway identification, only intact airways in cross-section (i.e. a complete circular structure) are identified. This potentially limits the number of airways detected, particularly in sections that have large amounts of tissue artefact due to histological tissue preparation. In this current work we have applied the model to mouse lung tissue. Further validation will explore its utility in human lung tissue.

In summary, we have presented a novel, powerful, semi-automated method that allows detailed, unbiased assessment of airway remodelling changes in the lungs of animals from *in vivo* models of asthma. The method has demonstrated important differences in the spatial remodelling of ECM, ASM and the epithelial layer in airways of various sizes, and key differences in the resolution of such remodelling changes after cessation of airway challenge.

## Acknowledgements

This study was funded by the Medical Research Council (MRC), Grant Number MR/M004643/1.

