## Supplemental Material for "Differential remodelling in small and large murine airways revealed by novel whole lung airway analysis"

**Supplementary methods and results**

Amanda L Tatler[1+], Christopher J Philp[1+], Michael R Hill[2+], Sam Cox[3], Andrew Bullock[2], Anthony Habgood[1], Alison John[1], Robert Middlewick[1], Katherine E Stephenson[1], Amanda T Goodwin[1], Charlotte K Billington[1], Reuben D O'Dea[2], Bindi S Brook[2♯], Simon R Johnson[1♯*]

1. Centre for Respiratory Research, NIHR Biomedical Research Centre and Biodiscovery Institute, School of Medicine, University of Nottingham.
2. School of Mathematical Sciences, University of Nottingham.
3. Digital Research Service, University of Nottingham.

**+ First authors contributed equally to this manuscript**
**# Joint senior author**

**Methods**

*Ethical statement for animal use*

All animal work was approved by the Animal Welfare and Ethical Review Board (AWERB) of the University of Nottingham (UK) and conducted in accordance with all terms of the Establishment, Project and Personal Licenses issued by the Secretary of State for the Home Office. Consistent with all national and international law, studies were carried out as detailed in the Animal [Scientific Procedures] Act 1986 (Amended Regulations 2012) (ASPA), Animal Welfare Act 2006, Directive 2010/63/EU, the LASA guidelines and in respect to the principals of Replacement, Reduction and Refinement. Work was performed under license number RGJ 40/3709 using the 19b5 protocol.

*Animals and the Ovalbumin Model of Asthma*

Female BALB/C mice aged 5-6 weeks were obtained from Harlan Laboratories Ltd. (UK) Animals were housed in a Specific Pathogen Free (SPF) environment in the Bio Support Unit, University of Nottingham, UK. Mice were randomly divided into a maximum of 4 mice per cage. Cages used were Techniplast GR500 Individually Vented Cages (IVC) at 21˚C and 50% humidity with environmental enrichment. Bedding consisted of Nestpak grade 5 sawdust (Datesand Ltd., UK) and Sizzle-Nest (Scanbur, Denmark). Mice were fed *ad libitum* 2018S Teklad Global 18% Protein Rodent Diet (Harlan Laboratories, UK) with constant access to fresh, filtered water and maintained on a 12-hour light, 12-hour dark cycle. Food and bedding were sterilized by autoclaving prior to use. Following receipt, animals were left in their new environment for 7 days to acclimatize before any experimental work began. Animals underwent IP sensitization to ovalbumin (OVA) and subsequent orophanarangeal challenge with OVA under anaesthetic (AB-G) of inhaled, 1% isofluorane in oxygen as previously published[1]. Samples were collected as described below with the first sample (day 34) collected 24 hours after the final inhalation challenge.

*Sample and Tissue Preparation*

At days 34, 35, 37, 39 and 41, animals were sacrificed by terminal anaesthesia (AC) (Euthatal – Merial Animal Health Ltd., UK) and confirmed with the absence of reflex motion. Sacrifice was conducted in accordance with ASPA Schedule 1. Bronchioalveolar lavage (BAL) samples were collected from each animal by tracheal cannulation and flushing of the lungs with 1ml sterile PBS and collected in Eppendorf tubes for further processing. After BAL collection, lung vasculature was perfused by transcardial perfusion with 40UI/ml heparin in normal saline to remove blood from the lung. Following this, lungs were inflated at 20cm/$H_2O$ with 4% formaldehyde. One inflated, the trachea was tied closed, lungs were excised with the heart attached and placed in 4% formaldehyde for 24 hours at room temperature. The whole respiratory tract was sectioned along the coronal plane, butterflied and paraffin wax embedded.

*Bronchioalveolar Lavage Preparation and Staining*

BAL samples were kept on ice until processing. All samples were spun at 1,500rpm at using an Eppendorf 5415D centrifuge for 5 minutes. Supernatant was collected and the pellet resuspended in 1ml PBS. Once resuspended the samples were spun again at 1,500rpm for a further 5 minutes. Supernatant was removed and the pellet resuspended in 500μl PBS. 200μl of each clean BAL sample was loaded into Shandon™ Cytofunnels (Thermo Scientific, UK) prepared with filter paper and Superfrost Plus adhesion microscope slides (Thermo Scientific, UK). Cassettes were spun at 450rpm for 6 minutes in a Shandon™ Cytospin 3 (Thermo Scientific, UK). Slides were removed from the cassettes and left to air dry for 15 minutes before staining.

Cytospin slides were stained using the Rapi-Diff II kit (Atom Scientific, UK). Slides were immersed in fixative solution A for 10 seconds. They were then transferred, without wiping or rinsing, into solution B. All slides were dipped 5 times into the solution while allowing 2 seconds between each dip. Excess stain was then blotted off the corner of the slide and the slide rinsed with PBS. Slides were then transferred to solution C and treated exactly as in solution B. Excess solution was rinsed from the slides in running tap water then the slides were blotted dry from the corners and left to air dry before examination under the microscope.

Cells were counted under a Nikon Eclipse 90i inverted light microscope (Nikon Instruments Europe B.V., UK) by a user blinded to animal treatment. Counting was performed based on cell staining and morphology and presented as percentage of total cells.

*Immunohistochemistry*

Paraffin wax-embedded tissue was deparaffinised using Histoclear (National Diagnostics) and rehydrated in a graded alcohol series. No antigen retrieval was performed. Endogenous peroxidases were quenched using 1% hydrogen peroxide in $dH_20$ for ten minutes. Slides were washed 3 times in PBS before blocking with 2.5% Normal Horse Serum (Vector Laboratories, CA, US). Anti-alpha smooth muscle actin primary antibody (ab5694) was diluted in blocking agent to 0.05μg/ml and the slides incubated in its presence overnight at 4°C. The Impress Anti-Rabbit secondary antibody kit (Vector laboratories, CA, US) was used according to the manufacturer's instructions. Colour was developed using DAB Peroxidase (HRP) Substrate Kit (Vector Laboratories) and counterstained with Haematoxylin.

*Picro-sirius Red Staining*

Picro-Sirius Red (Abcam, UK.) was applied to deparaffinised slides for 60 minutes before being rinsed in 2 changes of 5% acetic acid. These were subsequently rinsed in absolute ethanol before being dehydrated in 2 further changes of absolute ethanol. All slides were mounted with a glass cover slip and Vectamount permanent mounting medium (Vector Laboratories, CA, US).

*Airway Identification and Analysis*

The slides with histological sections stained for alpha airway smooth muscle actin (SMA) and collagen with picrosirius red (PSR) were imaged with a Hamamatsu Digital Nanozoomer 2.0-HT (Hamamatsu Photonics, UK). The *.ndpi images were converted to *.tiff format using ndpi2tiff (https://www.imnc.in2p3.fr/pagesperso/deroulers/software/ndpitools/) at maximum resolution and imported into MATLAB (The MathWorks Inc.) for image processing (the image processing code is available here: https://github.com/BindiBrook/AirwayIdentification). Two reference tiffs (one stained with aSMA and the other with PSR) for demonstration purposes are provided here: reference images.

Custom scripts were written to iterate through the images for processing and subsequently to measure the area of stained protein. Initial image processing comprised two steps. First, a pre-processor was used to find potential airways by locating approximately circular shapes. Next, a filtering process was used to retain objects with characteristics associated with airways, and eliminate the remainder. This second step was necessary because several hundred potential airways were typically identified by the pre-processor. Airway composition was determined by thresholding of image data; pixels within threshold values were deemed to represent stained protein area. Slightly different approaches were required for airway smooth muscle actin (SMA) and collagen (from the PSR stain) images, and in each case, a user-interactive process allowed for verification or manual adjustment. Full details of the image processing and analysis scripts are given below.

***Airway Identification:*** A custom preprocessor MATLAB program was developed to identify potential airways within the Nanozoom image; each such object is saved to a separate image file, together with their global location. Irregular circular objects were identified via a multi-step process performed on each image. Each image was converted to grayscale, and the contrast was adjusted using MATLAB functions *imadjust* and *stretchlim*; the result was binarized via *imbinarize*. Breaks in the epithelium of potential airways were closed via the *imerode* function and the space in the lumen of the object was filled via *imfill* (this applied to the complement of the eroded image). To determine the lumen area of these potential airways, lumen perimeter was smoothed via *bwmorph* and any stray pixels resulting from this procedure filled with *imfill*; the resulting object is labelled and its area determined with *bwlabel*.

The resulting set of potential airways is reduced by setting minimum (25$\mu$m) and maximum (500 $\mu$m) effective diameters (computed as the diameter of a circle of equal area), with objects with diameter outside the selected range being discarded. The boundaries of the retained objects (*i.e.* the potential airway lumen) were traced with *bwtraceboundary* and the ratio of the object area to its perimeter was computed. This metric was employed to eliminate highly convoluted objects; lower limits on this value were set by trial-and-error and objects with ratio exceeding the lower limit were discarded. For ASM images the lower limit was set as 21.25 for SMA images and and for PSR images this lower limit was 40.

The set of objects resulting from the initial selection process is still typically large and so a custom filtering algorithm, was used to retain objects most likely to be airways by calculating three separate metrics: ‘theta’, ‘transitions’ and ‘density’. These seek to capture the typically more regular shapes and thicker wall associated with airways, in contrast to (e.g.) alveoli. *Theta* measures the mean angle between a normal vector to the boundary trace defined at a given boundary point, and a line connecting the boundary point and the object centroid. *Transitions* measures the number of zero crossings in the binarized image along the length of an outward normal vector to the boundary trace. The *density* of the (potential) airway wall was computed by dilating the lumen boundary by 40$\mu$m and computing the ratio of the sum of the greyscale 8-bit pixel intensity and the total number of pixels between the original and dilated boundaries. See Supplementary Figure 1.

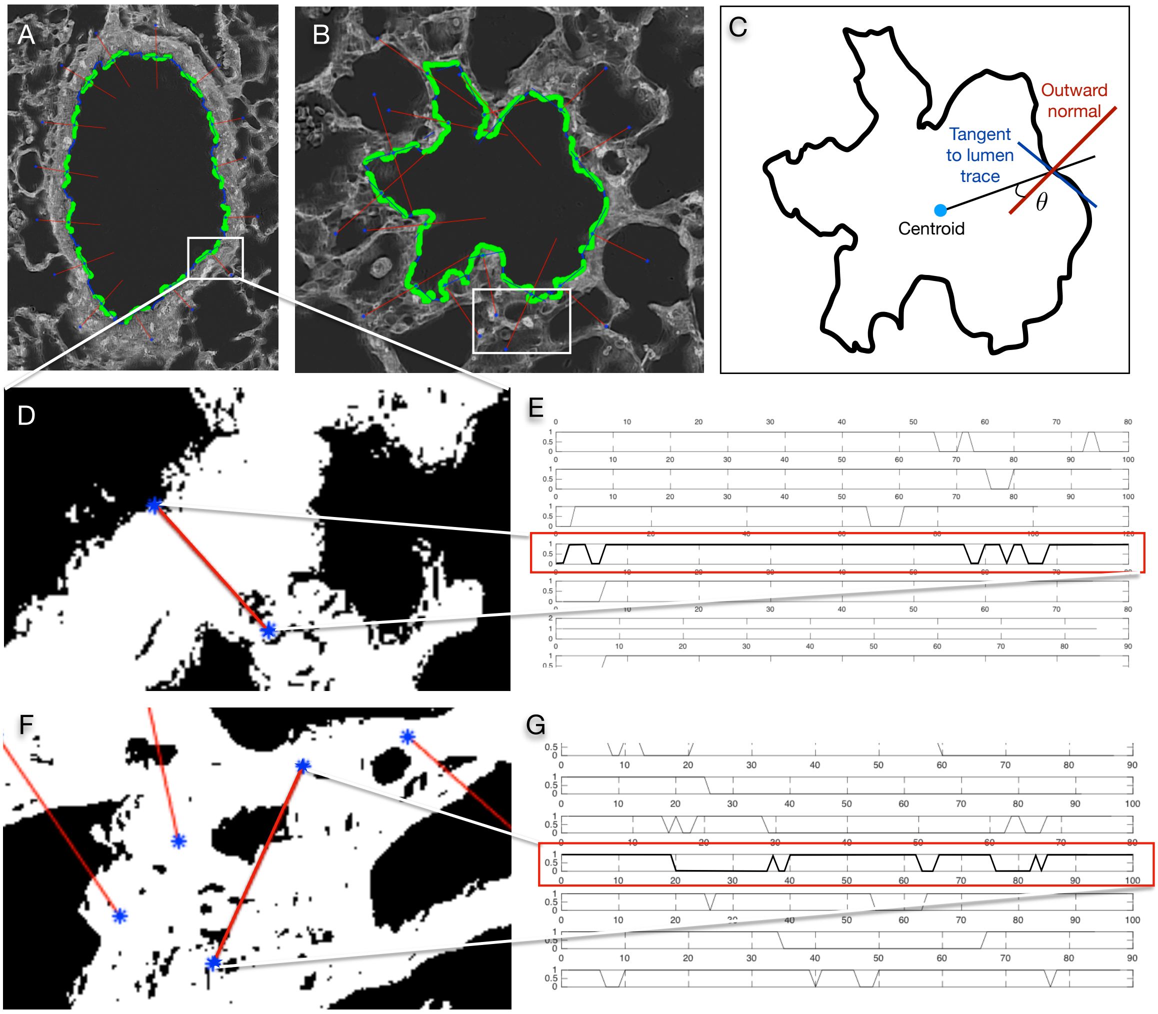


*Supplementary Figure 1:* ***Custom filtering metrics*** *‘Theta’ and ‘Transitions’ are based on 15 equally-spaced normal vectors to the boundary trace. Panels A and B show examples in a representative airway and non-airway object, respectively; a tangent, constructed from the green section of the boundary trace, is shown in blue; the normal is shown in red. The length of normal vector exterior to the boundary (marked by blue asterisks) is 100 pixels in each case. The circularity of identified objects is measured by an angle ‘Theta’ between the outward normal and a line connecting the object centroid to the normal (Panel C). Transitions captures multiple airspace crossings and internal wall structure associated with airways (Panels D,E) vs. alveoli (F,G) via the number of zero crossings in the binarized image along the length of the normal vector.*

For each potential airway, mean values of *transitions* and *theta* were computed from normal vectors of length 100 pixels defined at 15 equally-spaced boundary points. The following ranges were adopted for these three metrics depending on whether the image was SMA (or PSR): 0<*transitions*<501 (0<*transitions*<90), 0<*theta*<±111° (0<*theta*<±90°), 75<*density*<125 (83<*density*<141). A custom script is used to iterate through the filtered images for a final user-interactive visual identification of objects that are definitely airways, which are saved in a separate folder.

The suitability of the filtering thresholds was determined by testing on two manually-inspected Nanozoom images, for which the total number of true airways was known. The observed values for the metrics in these images were as follows (data for the subset of objects confirmed to be airways are given in brackets). *Transitions*: 54.6±35.7 on the range [0,501] (19.1±16.2 on range [0,90]); *theta*: 54.4±17.5 on the range [7.4,105.0] (31.1±13.6 on range [7.4,63.8]); *density*: 85.0±14.8 on the range [60.6,179.6] (110.2±7.4 on range [83.4,124.3]).

***Airway analysis:*** The script *main.m* is used to analyse the images identified as definitely airways for airway composition. To this end, in each airway image two further boundaries are defined in addition to the lumen trace described above. First, the user manually traced the outer boundary of the epithelium, representing the basement membrane (Supplementary Figure 2, blue line), and then a second boundary is automatically generated by dilating the region bounded by the basement membrane by 40$\mu$m (Supplementary Figure 2, red line).

To determine the area of stained SMA between the two boundaries defined above, the RGB values of each pixel in the region were converted to grayscale, and then the complement was taken. Pixels within a user-specified threshold intensity window (set via an interactive slider, so that the selected pixels can be visualised) were defined as representing SMA-stained tissue. Collagen-stained pixels are isolated by converting RGB values to CIE 1976 L*a*b* values using *rgb2lab*. Similar to the SMA case, the pixel threshold is set via a slider (to adjust the 'a' image component). In both cases, further manual adjustment allowed the user to erase and manually select pixels that s/he determined were incorrectly selected or omitted in either the grayscale (for SMA) or the L*a*b* (for PSR) threshold process.

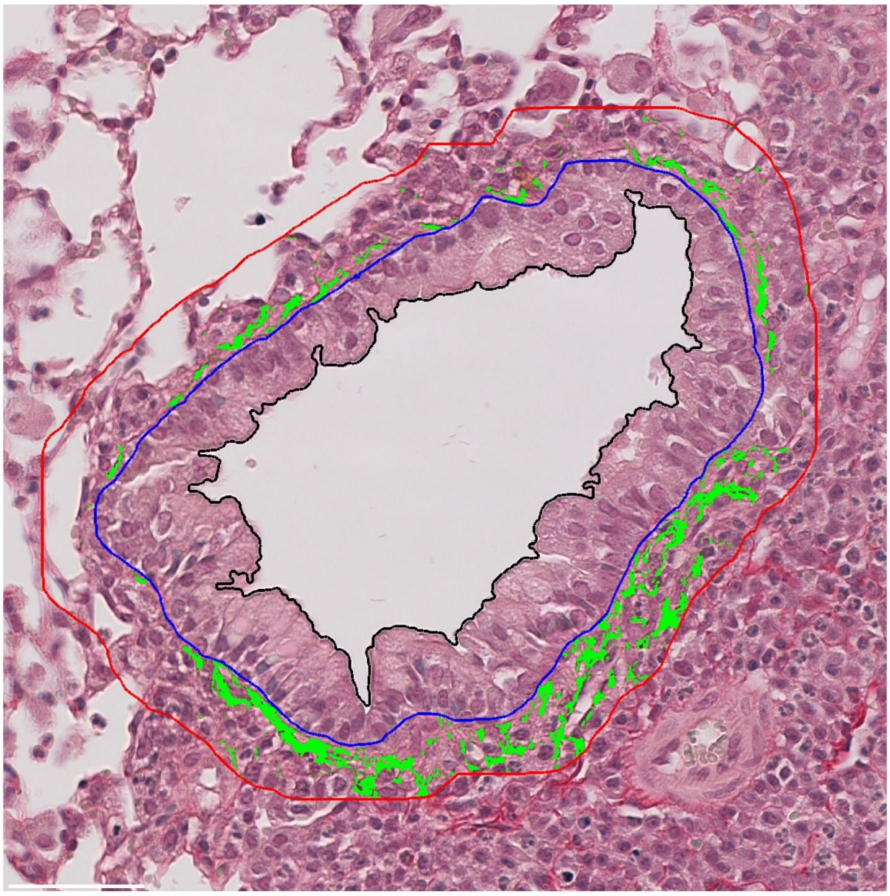

*Supp. Figure 2.* ***Airway composition analysis.*** *Example PSR-stained airway, showing the manually-traced basement membrane (blue line), and automatically-generated second boundary (red line) demarking the 40μm airway wall thickness. Pixels identified as PSR-stained tissue via a user-specified threshold intensity window are shown in green.*

*Data Analysis, Statistics and Graphical Software*

Data analysis and statistics were performed using custom scripts in MATLAB.

*Differences between individual animals and group sizes do not affect global remodeling data interpretation.* As described in the main text, we identify a total of 1007 PSR-stained and 1022 SMA-stained airways, comprising 90-240 airways (denoted $n_a$) from 4-10 mice (denoted $n_m$). To analyse changes in area fraction of ECM and ASM, we calculate: (i) the mean area fraction per mouse ($\mu_m$), (ii) the group mean across all the $n_a$ airways at each timepoint ($\mu_a$), and (iii) the mean for each group of animals ($\mu^*_m$); see Supplementary Table 1. We observe similar values for $\mu_m$ and $\mu^*_m$ across all groups and timepoints, and no significant difference between values of $\mu^*_m$ and corresponding $\mu_a$ (all p-values > 0.7; see Supplementary table 1). Moreover, one-way ANOVA shows that the global changes in ECM and ASM area fraction airway observed during the resolution period that are described in the main text for pooled data (with mean, $\mu_a$) persist when considering individual mice ($\mu_m$), though with somewhat weaker significance due to the reduced number of datapoints; see Supplementary Figure 3.

| ECM | D34 Control | | D34 Ova | | D41 Control | | D41 Ova | |
|---|---|---|---|---|---|---|---|---|
| | Means | n | Means | n | Means | n | Means | n |
| Pooled mean area fraction ($\mu_a$) | 0.1084 | 121 | 0.1654 | 110 | 0.1407 | 155 | 0.1901 | 124 |
| Means per mouse ($\mu_m$) | 0.1154<br>0.0674<br>0.0878<br>0.1131<br>0.1571<br>0.1571 | 12<br>37<br>21<br>14<br>10<br>27 | 0.1509<br>0.1842<br>0.1490<br>0.1533<br>0.1536<br>0.1935 | 14<br>16<br>23<br>18<br>15<br>24 | 0.1694<br>0.1253<br>0.1568<br>0.1231<br>0.1518<br>0.1389 | 25<br>29<br>18<br>39<br>16<br>28 | 0.1920<br>0.1533<br>0.1965<br>0.2296 | 34<br>25<br>50<br>14 |
| Mean based on number of mice ($\mu_m$*) | 0.1163 | 6 | 0.1641 | 6 | 0.1442 | 6 | 0.1928 | 4 |
| p values (cf $\mu_a$, $\mu_m$*) | 0.7184 | | 0.9580 | | 0.8742 | | 0.9264 | |

| ASM | D34 Control | | D34 Ova | | D41 Control | | D41 Ova | |
|---|---|---|---|---|---|---|---|---|
| | Means | n | Means | n | Means | n | Means | n |
| Pooled mean area fraction ($\mu_a$) | 0.0754 | 164 | 0.1346 | 158 | 0.0785 | 138 | 0.1028 | 90 |
| Means per mouse ($\mu_m$) | 0.0486<br>0.0810<br>0.1103<br>0.0408<br>0.0559<br>0.0991 | 28<br>51<br>17<br>17<br>20<br>31 | 0.1264<br>0.1338<br>0.1464<br>0.0585<br>0.1251<br>0.1867 | 20<br>16<br>34<br>31<br>26<br>31 | 0.1116<br>0.0688<br>0.0947<br>0.0529<br>0.0424<br>0.1068<br>0.0599<br>0.1223<br>0.1343<br>0.0453 | 10<br>16<br>15<br>12<br>14<br>9<br>12<br>22<br>5<br>23 | 0.0991<br>0.1502<br>0.0964<br>0.0571 | 14<br>19<br>44<br>13 |
| Mean based on number of mice ($\mu_m$*) | 0.0726 | 6 | 0.1295 | 6 | 0.0839 | 10 | 0.1007 | 4 |
| p values (cf $\mu_a$, $\mu_m$*) | 0.9215 | | 0.8812 | | 0.7782 | | 0.9448 | |

*Supp. Table 1. **Global ECM and ASM area fractions during resolution period.** Pooled mean ASM and ECM area fraction across all airways ($\mu_a$), mean area fraction per mouse ($\mu_m$) and the mean for each group of animals ($\mu^*_m$) over the resolution period day 34 to 41. Group means ($\mu^*_m$) do not differ significantly from pooled data ($\mu_a$) as indicated by p-values obtained from one-way ANOVA; further statistical analysis of ECM and ASM data during the resolution period is presented in Supplementary Figure 3.*

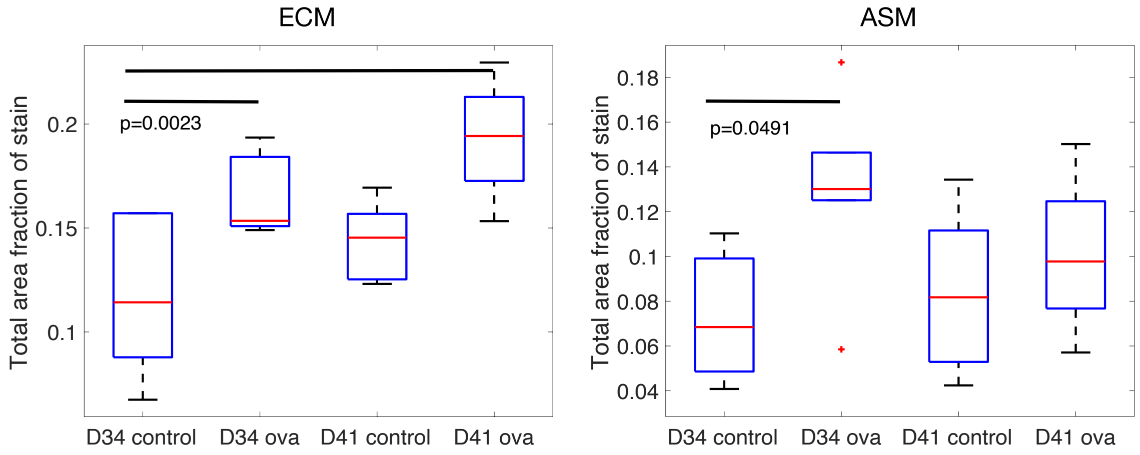


*Supp. Figure 3: **Global changes in airway constituents during resolution period, based on data grouped per mouse.** Box plots showing change in area fraction $\mu_m$ of ECM (left) and ASM (right) from days 34 to 41 (full data presented in Supplementary Table 1). On each box, the central mark indicates the median, and the bottom and top edges of the box indicate the 25th and 75th percentiles, respectively. The whiskers extend to the most extreme data points not considered outliers. Solid black horizontal bars and given p-values indicate means that are significantly different to mean at day 34 control (determined via one-way ANOVA).*

Intrasubject heterogeneity is exposed by analysis of the full set of airways. In (e.g.) Supplementary Table 1 and Supplementary Figure 3, a rich dataset is collapsed into a single metric. Supplementary Figure 4 exemplifies the dataset heterogeneity within each mouse, showing the ECM remodelling data (ECM area fraction, basement membrane perimeter and radial variation in ECM content) at day 34 within control and OVA animals. (Similar data is observed for ASM remodelling, but is not shown here for brevity.) For example, the distribution of total ECM datapoints in each mouse varies widely, in both control and OVA conditions; wider spread in each mouse but somewhat more consistency between mice is observed in the OVA animals (Blue lines in row 1 in panels A and B, Supplementary Figure 4). Similar features are observed in the radial variation of ECM deposition (row 3 in Panels A, B, Supplementary Figure 4); interestingly, however, we observe that the distributions of basement membrane perimeter show somewhat more consistency between animals than those for ECM remodelling (cf. rows 1 and 2 in Panels A and B, Supplementary Figure 4), thereby underscoring the significant intra- and inter-subject heterogeneity in airway constituent remodelling. The intra-subject heterogeneities are exposed in the airway size-based data stratification that we exploit to interrogate differential remodelling in the main text.

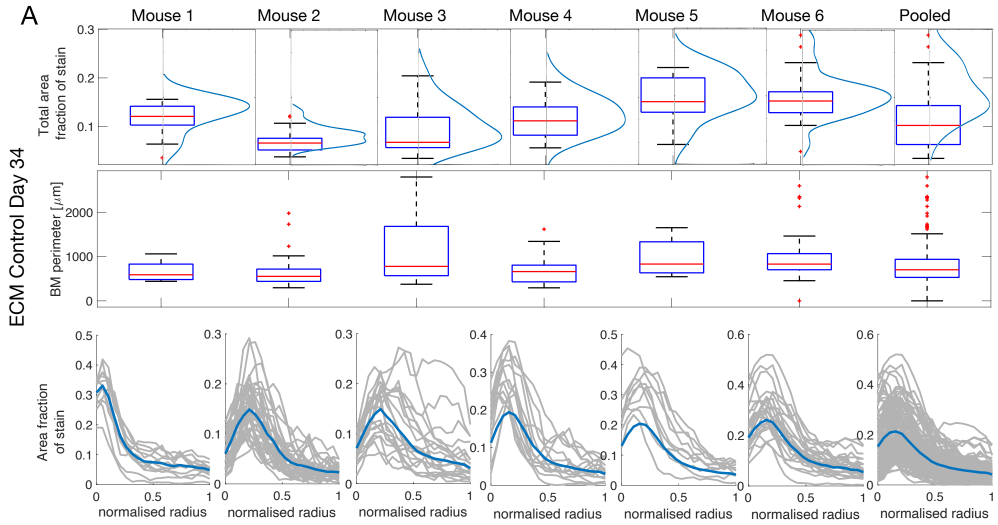

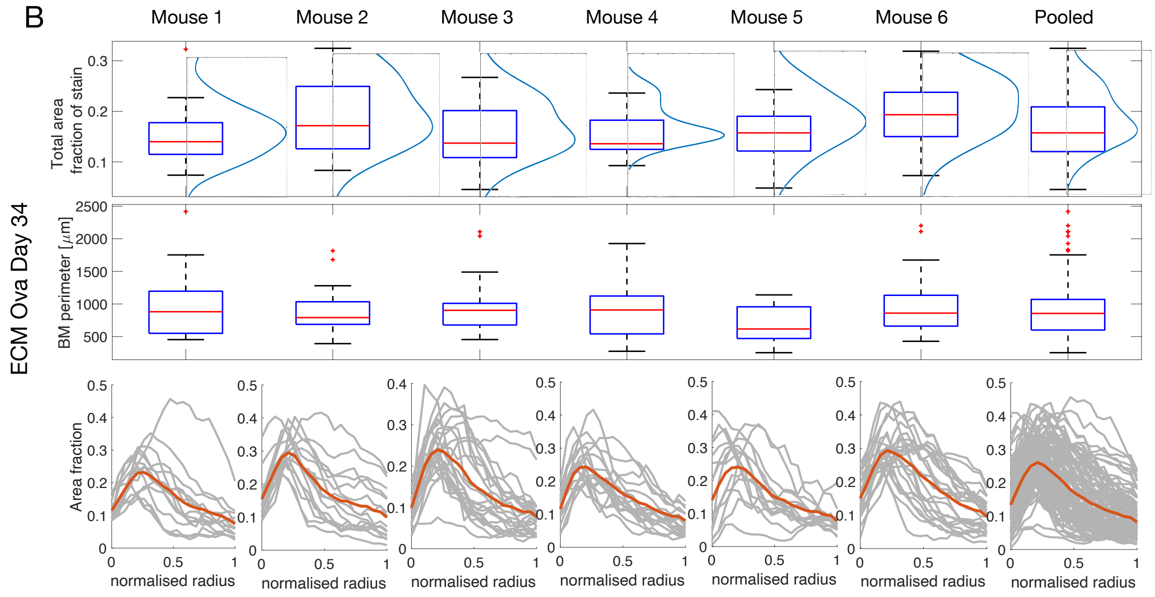


*Supp. Figure 4:* ***Intrasubject ECM remodeling heterogeneity at day 34****. ECM remodeling in control (A) and ova (B) animals. Row 1: Box plots showing ECM area fraction in each mouse, and in the pooled dataset. The central mark indicates the median, and the bottom and top edges of the box indicate the 25th and 75th percentiles, respectively; the whiskers extend to the most extreme data points not considered outliers. The data distribution is indicated by the blue curves in each case, [fitted via kernel smoothing functions]. Row 2: Box plots showing the basement membrane (BM) perimeter in each mouse. Row 3: Spatial distribution of ECM area fraction (computed as explained in the main text) across the radial thickness of the airway wall, normalized to [0,1].*

*Fine-grained stratification of airway remodelling by airway size reveals differential remodelling dynamics.* Supplementary Figure 5 provides 2D projections of 3D histograms to highlight how the frequency of airways of given ASM/ECM airway fraction and airway size (as defined by basement membrane (BM) perimeter) evolves during the remodelling period.

Panels (A-D) provide schematic diagrams to aid interpretation of the panels (E-H), as well as Fig 4 in the main text. Panel (A) represents control data at day 34; if all airway sizes (quantified by BM perimeter) displayed identical area fraction increase over the OVA challenge period, while remaining of the same size, the pixels in (A) would shift horizontally to a larger area fraction as shown in (B). The best-fit line through this data (solid blue line in panel (B)) would hence be parallel to that in (A) (here, the best-fit line from A is shown as a dashed line for comparison). If, on the other hand, larger airways remodelled more than smaller ones (with airway sizes remaining fixed), the horizontal shift would be larger for larger values of BM perimeter and the best-fit line would have smaller slope (panel (C)). Alternatively, if the size of the larger airways increased, but area fraction remained constant, the slope would increase, as indicated by the shift in panel (D).

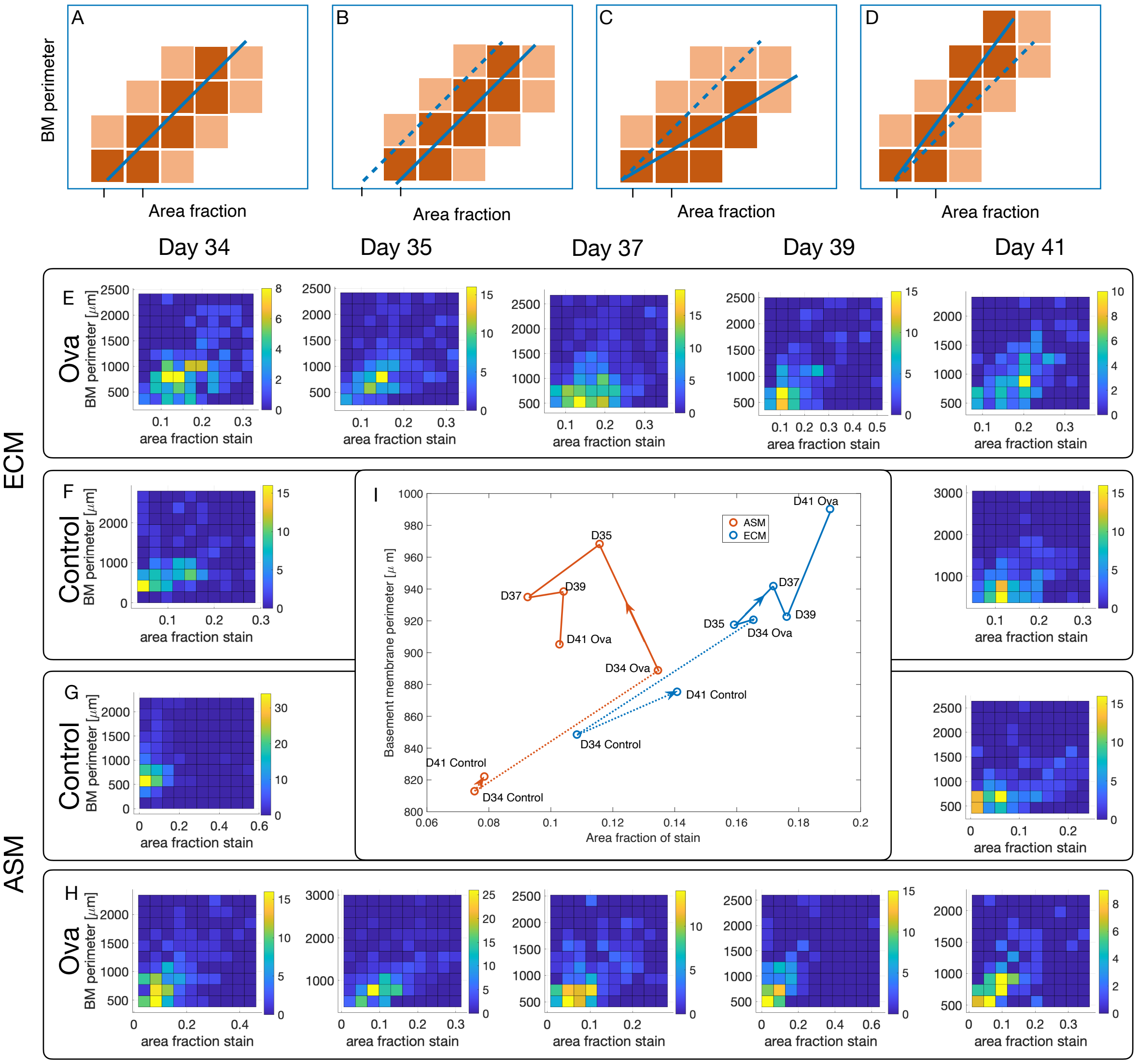


*Supp Figure 5:* ***Fine-grained stratification of remodeling data by basement membrane perimeter.*** *Data is presented as 2D projections of 3D histograms. Colour intensity indicates the frequency (see colour bars) of an airway of given basement membrane (BM) perimeter and ASM/ECM area fraction occurring in the control/ova datasets. Panels (A-D): Schematic diagrams to aid interpretation of the remaining panels; solid blue lines are lines of best fit through the data; the dashed blue line in B-D is that corresponding to A, included for comparison. Panels (E,F) show ECM data over the resolution period at days 34-41 (OVA) and at day 34, 41 (Control); (G,H) show corresponding histograms for ASM remodeling. Panel I shows the position of the mean (in each coordinate direction) of the data shown in panels (E-H) to highlight the differential resolution dynamics.*

Panels (E-I) show the differential remodelling with airway size described in the main text, and the strong temporal variation over the resolution period. In particular, panel (I) highlights the resolution of the ASM almost to baseline, while ECM remodelling continues through this period.

*Mean lumen area was not significantly altered in ovalbumin-challenged compared to control animals.* As described in the main text, somewhat surprisingly, the expected decrease in lumen area associated with increased epithelial area was not observed (Figure 6, 7, main text). This can be explained via geometrical arguments and simulated data. Supplementary Figure 6A indicates the relevant measures obtained from image data that we refer to below.

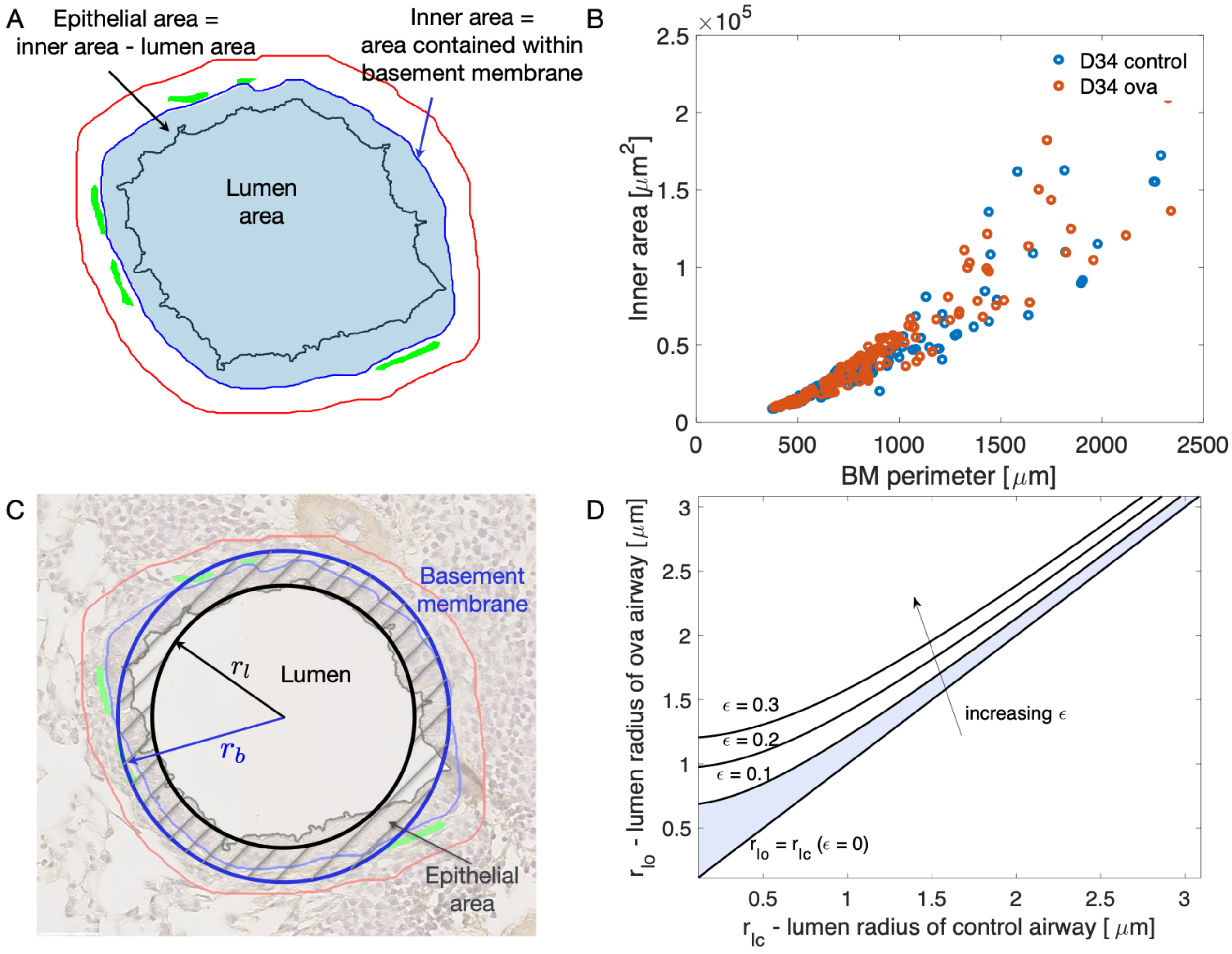


*Supp. Figure 6:* **Epithelial involvement in remodeling.** *(A) Schematic diagram highlighting the three independent measurements that are made from image data of each airway: BM perimeter, inner area and lumen area. Epithelial area changes reported in the main text (Fig 6, main text) are determined by subtracting the lumen area from the inner area. (B) BM perimeter and inner area data for all airways at day 34. (C,D) Geometrical arguments and simulated data to explain the relationship between epithelial area and BM perimeter observed in the main text (Figs 6, 7, main text). (C) Schematic diagram to indicate the equivalent circles employed to represent the lumen boundary and basement membrane. (D) Lumen radius of a control airway ($r_{lc}$) and an that of an OVA airway ($r_{lo}$), for simulated data obtained according to equation (S2), where the control BM radius is fixed at $r_{bc} = 4.5$). The shaded region indicates values for which the inequality (S2) is satisfied (when $\epsilon = 0.1$) and the lumen area in OVA airways exceeds that in control animals.*

Approximating the basement membrane and lumen by circles (Supplementary Figure 6C) with radii $r_b$ and $r_l$, the epithelial area, normalised by basement membrane perimeter, can be computed as

$$E = \frac{r_b^2 - r_l^2}{2r_b}. \qquad (S1)$$

Now, introducing an additional subscript to distinguish between OVA challenge ($E_o, r_{bo}, r_{lo}$) and control ($E_c, r_{bc}, r_{bl}$), and assuming that OVA challenge leads to a small increase $\epsilon$ in BM radius over control conditions ($r_{bo} = r_{bc} + \epsilon$) the normalised epithelial area in each case can be computed via equation (S1). Demanding that the epithelial area *and* lumen area in OVA challenge is greater than that in control (*i.e.* $E_o > E_c$, $r_{lo} > r_{lc}$) then leads to the following inequality:

$$r_{lc} < r_{lo} < \left[ r_{lc}^2 + \epsilon \frac{r_{bc}^2 + r_{lc}^2}{2r_{bc}} + \epsilon^2 \right]^{1/2}. \qquad (S2)$$

Given that we have observed that the basement membrane perimeter is not significantly changed by OVA challenge (see main text, and Supplementary Figure 6B highlighting that BM perimeter and inner area are highly correlated in OVA and control conditions, as expected), equation (S2) provides a condition on the lumen radius that allows for both epithelial and lumen area in OVA animals to exceed those in control conditions.

Supplementary Figure 6D shows the relationship between $r_{lo}$ and $r_{lc}$ (for fixed $r_{bc}$) and highlights the region in which the inequality (S2) is satisfied.

Figure 7 in the main text shows simulated OVA data derived from day 34 control data to illustrate the applicability of this theoretical result. Here, we compute simulated OVA data by adding a small random perturbation to paired (lumen area and inner area) control data at day 34 via $E_{lo} = E_{lc} + \xi_l$ and $E_{io} = E_{ic} + \xi_i$, where subscripts $i$ and $l$ denote inner and lumen areas respectively, $\xi \sim N(\mu, \sigma^2)$ are normally-distributed random numbers with mean $\mu$ and standard deviation $\sigma$; specifically $\xi_l \sim N(3,2)$ and $\xi_i \sim N(5,2)$, chosen to qualitatively match experimentally observed changes in the ova compared with control distributions. Corresponding epithelial area is computed from the difference of these areas.
